# Novel universal domain-centric method for protein classification

**DOI:** 10.1101/2025.03.28.645719

**Authors:** Shakiba Fadaei, Fanny S Krebs, Vincent Zoete

## Abstract

Human protein kinases constitute a large superfamily of about 500 genes, historically classified into subfamilies based on phylogenetic relationship. However, many kinases remain unclassified. Phylogeny is typically based on multiple sequence alignments, and neglects the physico-chemical properties of residues at each position of the sequence. By incorporating these properties, we can gain deeper insights beyond basic alignments. Here we use, for the first time, a detailed physico-chemical description of kinases to identify class-specific structural regions, supporting an unsupervised classification method capable of classifying previously unlabeled kinases. This novel approach aligns with existing phylogeny-based classifications while offering refinements and enhanced accuracy. Ultimately, we use machine learning techniques to classify unlabeled kinases, validated by analyzing class-specific structural regions. This new classification approach goes beyond current rankings and can be applied to any type of protein, such as immunoglobulins and G protein-coupled receptors.

## Introduction

Protein phosphorylation is a crucial regulatory process [1,2], with kinases playing a key role in orchestrating it to modulate cellular functions and signaling pathways [3,4]. Kinases activity imparts a vast functional diversity of proteins, enabling them to carry out a variety of specialized tasks and respond dynamically to cellular signals [5]. For example, cell growth, proliferation, differentiation, protection, and degradation are all regulated by protein kinases [6–9]. Given the prominent role of kinases in every important aspect of a cell’s life, their implication in diseases is not surprising [10–13]. The cataloging of the human kinome began with the completion of the human genome project [13,14], which provided a foundation for identifying and characterizing various gene families, including protein kinases [15–19]. With the near completion of the human genome in the early 2000s, a first catalog of human kinases was compiled. Manning et al. employed a hidden Markov model (HMM) profile of protein kinases to identify and classify additional human protein kinases [20]. They found 518 human kinases, including 40 atypical protein kinases, which have biochemical kinase activity but low sequence similarity to other kinases, and classified the typical kinases into seven classes: AGC, CAMK, CK1, CMGC, STE, TK, and TKL [20]. In a later work, a structure-based multiple sequence alignment (MSA) of 497 kinases was provided by the KinCoRe database [21–23].

Some newly identified kinases exhibit sequences that do not resemble those of any known classes [24]. Consequently, analyzing these kinases solely based on sequence similarities is insufficient. This highlights a gap in our molecular-level understanding of kinases [1]. Knowledge of protein 3D structures is crucial for elucidating and understanding the structure-activity relationship of proteins and comprehending how they interact with other molecules, such as substrates or drugs [25–27]. This insight is crucial for biological research in general and for drug design in particular [28–30]. Protein kinases are structurally fascinating, as they achieve diverse specificities and mechanisms within a shared molecular framework [18]. They all have similar folds, with highly conserved active sites, despite a substantial sequence diversity. Kinases have an N-terminal and a C-terminal lobe [31,32]. The N-terminal lobe consists of five *β*-sheet strands and one *α*-helix, called the C-helix. The C-terminal lobe is made of six *α*-helices and two *β*-sheets. The two lobes are connected by a hinge [33]. The hinge region is where the adenosine 5′-triphosphate (ATP) binding site is situated [12]. The ATP binding site contains a DFG motif and an HRD motif, which are highly conserved across all kinases [34] (Figure 1).

**Figure 1.**
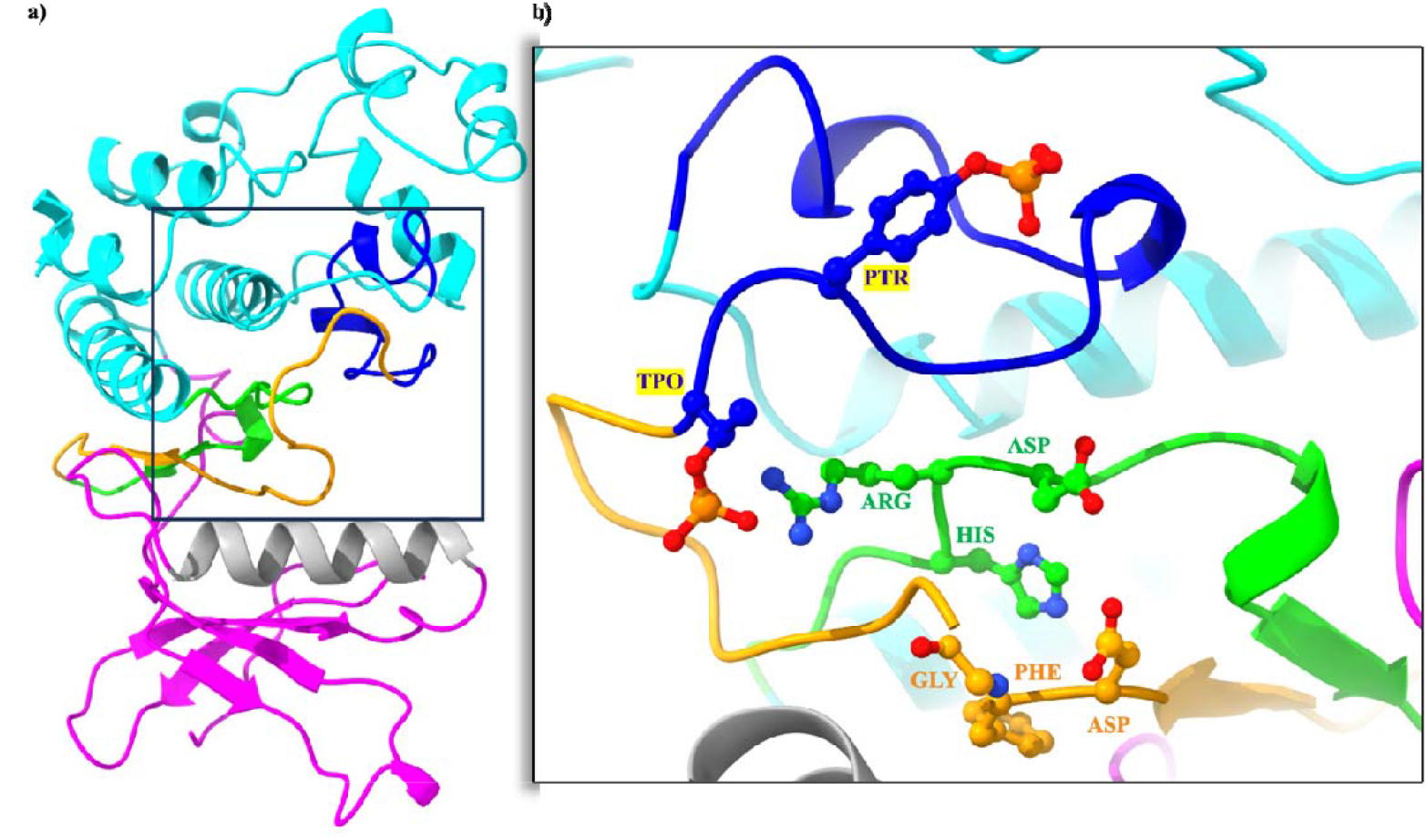
Structure of a typical protein kinase. **a)** Representation of the phosphorylated human MAPK13 kinase (PDB ID 4MYG [38]). The C-lobe is consisted of the cyan section; the dark blue section, which is the activation loop of the C-lobe; the catalytic loop colored in lime that contains the HRD motif. The N-lobe has - sheets in magenta; the C-helix in gray; and the activation loop of the N-lobe colored in orange, where the DFG motif is located. **b)** Shows the zoom-in section of the hinge, where the ATP-binding site is located. The residues that have the labels highlighted are post-translational modifications sites (PTM); PTR: phosphorylated tyrosine, and TPO: phosphorylated threonine. Residues constituting the HRD and DFG motifs are colored in lime and orange respectively.

Here, for the first time, we will exploit the KinCoRe [22] MSA and more importantly the physico-chemical properties of residues at each position of the kinase domain, to generate quantitative and unsupervised kinase classifications, and assess if they are consistent with phylogeny-based classifications [35]. This new approach can also be used to categorize kinases that were previously unclassified. Importantly, our new approach is universal, and could be applied without any change to other large families of proteins and protein domains, such as proteases, G protein-coupled receptor proteins or DNA-binding domains [36,37].

## Results

### Kinase physico-chemical representation and dimensionality reduction

Figure 2. shows the Uniform Manifold Approximation and Projection (UMAP) [39] visualization of the full-kinase representations consisting in a matrix of 2218 columns corresponding to multiple sequence alignment (MSA) positions and 30 rows corresponding to AAindex [40] for each sequence. The data points are color coded based on their class labels according to KinCoRe. Importantly, this classification was not utilized in the calculation and derivation of the PCA and UMAP analyses. Instead, it was solely employed to visualize the distribution of these classes within our strict AAindex-based UMAP representation. **Figure 2.a** reveals that the clusters obtained with our untransformed AAindex-based kinase representation are very compact, with a lot of overlap. Although two data points from the same class are very similar, they are not identical, which is due to data redundancy. To correct this, we performed a principal component analysis (PCA) of our kinase representation. The UMAP of the PCA-transformed kinases representation exhibits visible clusters of the same class (**Figure 2.b**), with less overlap and reduced compactness of clusters. For example, the data points of the “TYR” kinases, are more spread in **Figure 2.b**, which is the case for almost all classes. Importantly, the dispersion of the data points remains limited enough to preserve information about their main class, thereby leading data points of the same class to cluster.

### *k*-means unsupervised clustering *vs.* class labels

To determine the optimal number of clusters *k*, we fitted the PCA-transformed physico-chemical kinase representations for *k* values ranging from 2 to 30, and calculated the silhouette score (**Figure 3.a-i**). We observed local maxima for cluster numbers of 11, 14, and 18. The UMAP of the reference classes, and the unsupervised *k*-means clustering *k*=11, 14, and 18 clusters are shown in **SI Figure 1**.

**Figure 2.**
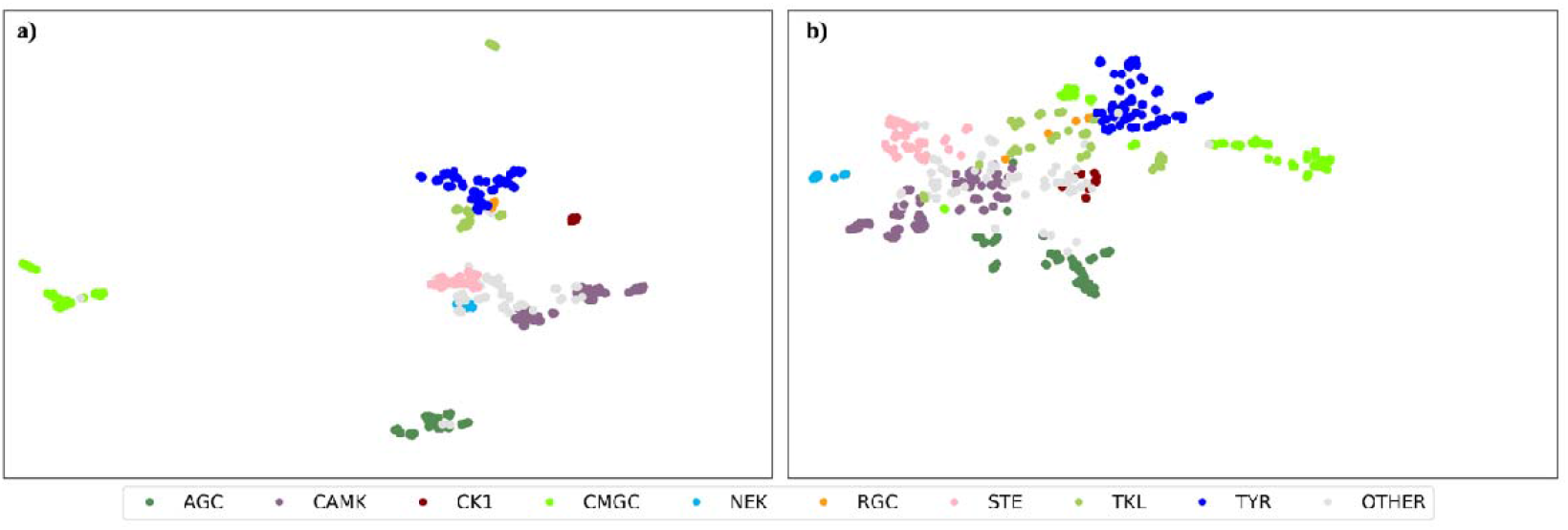
UMAP visualization color coded based on main class labels. **a)** UMAP visualization of the original kinase representation. **b)** UMAP visualization of the PCA-transformed kinase representations. Importantly, representation vectors do not contain any information about the classes.

**Figure 3.**
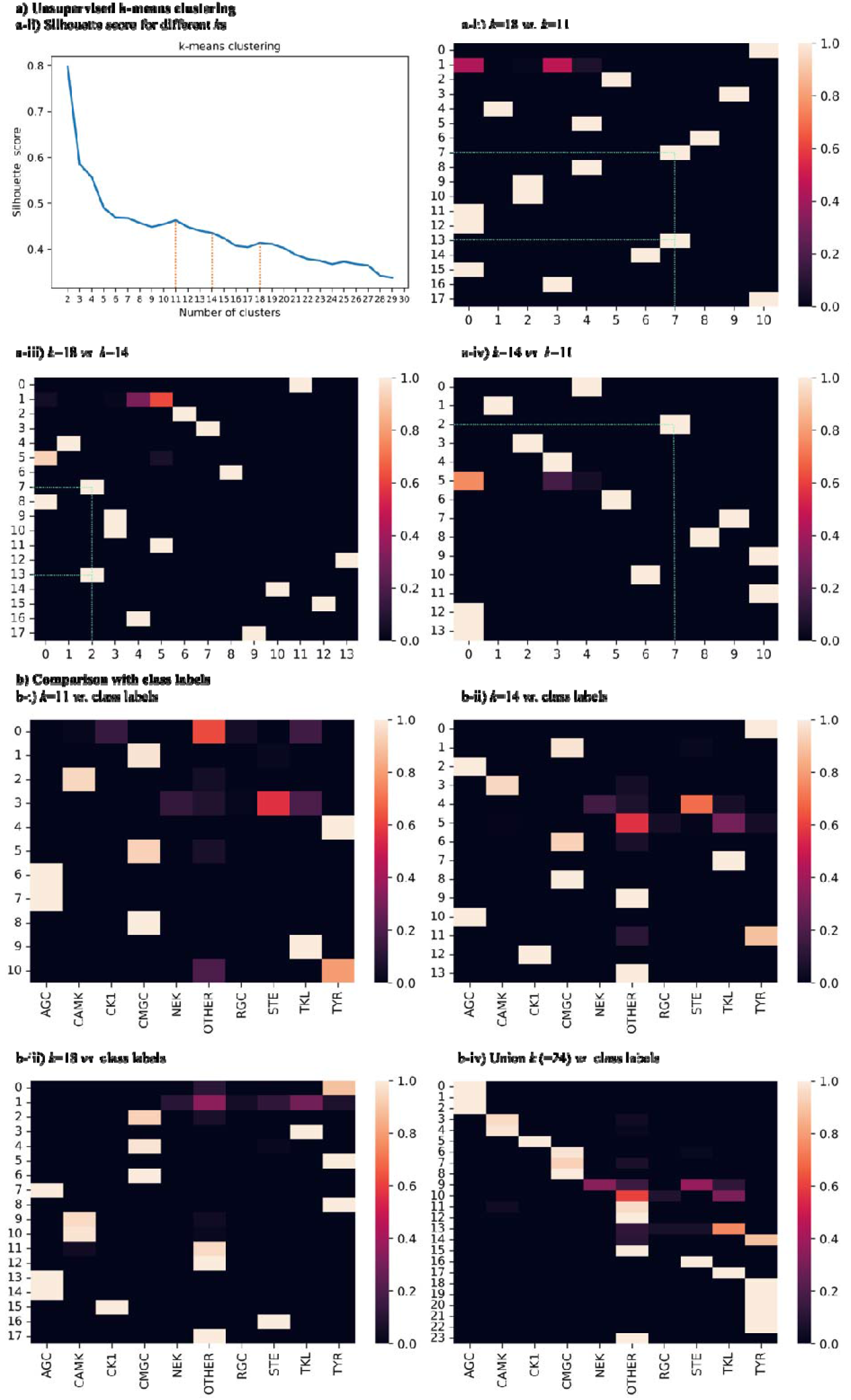
Unsupervised K-means clustering model. **a)** The clusters are assigned without any information about their class labels. **a-i)** The silhouette score plot for *k*s in the range of 2 to 30. The local maxima determine the most optimum number of clusters: *k*=11, 14, and 18. **a-ii)** Pairwise confusion matrix of cluster assignments k=18 vs. k=11. **a-iii)** Pairwise confusion matrix of cluster assignments *k*=18 *vs. k*=14. **a-iv)** Pairwise confusion matrix of cluster assignments *k*=14 *vs. k*=11. **b)** Comparing the k-means cluster assignments with the KinCoRe class labels. **b-i)** *k*=11 *vs.* class labels. **b-ii)** *k*=14 *vs.* class labels. **b-iii)** *k*=18 *vs.* class labels. **b-iv)** Union of all three cluster assignments (*k*=11, 14, and 18), resulting in 24 distinct clusters *vs.* class labels.

To investigate if the assigned clusters of *k*=11, 14, and 18 are consistent with one another, we represented the cluster assignments using pairwise confusion matrices (**Figure 3.a-ii,iii,iv**). The y-axis corresponds to the clustering with the higher number of clusters, and the x-axis corresponds to the lower ones. The heatmap is normalized based on rows—a square on the heatmap with a value of one means that all the data points in the corresponding cluster on the y-axis are also in one cluster shown on the x-axis. We will refer to subclusters as data points assigned to a single cluster at one *k* value but split into multiple clusters at another *k* value. For example, in **Figure 3.a-ii** the content of clusters 7 and 13 of *k*=18 are grouped into cluster 7 of *k*=11. **Figure 3.a-iii** shows light colored squares corresponding to cluster 2 of *k*=14 on the x-axis, and clusters 7 and 13 of *k*=18 on the y-axis. **Figure 3.a-iv** shows that cluster 2 for *k*=14 is identical to cluster 7 for *k*=11. The lighter-colored squares in the heatmaps indicate that most clusters are highly consistent and closely aligned. Meaning the same data points were clustered together for both *k*s. The clusters that might have inconsistencies are clusters 1 and 5 for *k*=18 and cluster 5 for *k*=14. We ultimately created an overall clustering based on the union of the different *k* values, where clusters were defined as containing data points that are present in the same combination of clusters for the *k*=11, 14, and 18 clustering. This led to the clustering containing 24 different clusters. (**SI Table 1**).

Clusters are considered consistent if the same data points are clustered together across different *k*s, i.e., if the corresponding data points were assigned to a given cluster at a low *k* value, they were either assigned to a single cluster or to multiple subclusters at a higher *k* value. On the contrary, they are considered inconsistent if the corresponding data points belong to one cluster at a high *k* value but are split into several clusters at a lower *k* value. Based on the analysis of consistent and inconsistent data, we established a confidence score for each group (**SI Table 1**). This scoring ranges from 5 for inconsistent clusters to 10 for totally consistent clusters.

364 data points are assigned to consistent clusters across different clustering algorithms. 16 out of the 24 clusters are consistent across different k values, with the following numbers in our new assignments: 0, 1, 2, 3, 4, 5, 6, 7, 8, 11, 12, 14, 16, 17, 19, and 23. 133 data points are assigned to inconsistent clusters, meaning they might belong to one cluster for *k*=18, but in two separate clusters for *k*=11. In our union cluster assignments, we placed these data points into two different clusters with a lower confidence score.

We measured the overlap of our defined clusters based on *k*-means clustering with their KinCoRe class labels. **Figure 3.b** shows the confusion matrix of each cluster number and the class labels for different *k*s. Based on this classification, our final unsupervised assigned clusters for *k*=24 overlap with labeled kinases for 90% of data points. Furthermore, we compared the union *k*=24 clusters with the kinases classes. Many clusters covered only one class, represented in light color (**Figure 3.b-iv**), and validated the performance and accuracy of our unsupervised classification. Based on the confidence score assigned (see **SI Table 1**), clusters 0, 1, 2, 3, 4, 5, 6, 7, 8, 11, 12, 14, 16, 17, 19, and 23 have the highest cluster quality in our different *k* cluster assignment, which is measured by coherence of data points clustered together for different *k*s. Clusters 0, and 1 are sub-clusters for the “AGC” class. Cluster 3 contains data points from “CAMK” and “OTHER”. Since cluster 4 has a high confidence score, it is interesting to further study if the kinases corresponding to “OTHER” data points in this cluster could be reclassified as “CAMK”. The “TYR” class is distributed across clusters 14 and 19. CMGC is in clusters 6, 7, and 8. Some clusters, such as 3, 4, 6, 7, and 14, contain data points from multiple classes, usually shared between “OTHER” and one other class. Again, such overlaps could be leveraged to further classify unlabeled kinases in an unsupervised manner. To compare the new unsupervised classifications to the KinCoRe ones, we calculated pairwise intra- and inter-cluster similarities for both methods (**SI Figure 2** and **SI Figure 3**). An ANOVA test was performed, to calculate the p-value and F-statistics for one group versus all the others (**SI Table 2** and **SI Table 3**). While both techniques consistently returned low p-values for the clusters, the union *k*-means clustering produces higher F-statistic values than the KinCoRe alignments, suggesting that it captures significant group differences.

Clusters 11 and 12 have higher confidence in our unsupervised clusters and they are classifying kinases from the OTHER class. We performed the randomization test on these clusters versus all the other kinases to find what sets them aside. For example in Cluster 12, positions 423, 939, and 2176 are statistically distinct with effect size higher than 2.5. Studying these positions can give us insights into functionality of these kinases. However, doing a detailed structural and functional study of the clusters is out of the scope of the current work.

### Revealing the difference between the classes

Each PC is a linear combination of the original kinase representations, with the fitted weights for each PC remaining consistent across all data points and classes. “CMGC” and “TYR” PC1 values are at both extremities of the plot and are separate from the other classes (**Figure 4.a**). In **Figure 4.b**, the “AGC” PC2 value is higher than other classes. We observe differences in the range of values across classes. To name a few class pairs with different PC1 and PC2 ranges: “CAMK” and “TYR”, “CMGC” and “STE”, and “TKL” and “CAMK”. Some classes share the same range of PC2, but have distinct ranges of PC1, like “TYR” and “CMGC”. This provides valuable insight on how each PC explains a portion of data distribution, highlighting the importance of multiple PCs for kinase classification. Moreover, it reveals that physico-chemical properties of residues at certain positions impact PC values differently, indicating class-specific positions in the sequence.

**Figure 4.**
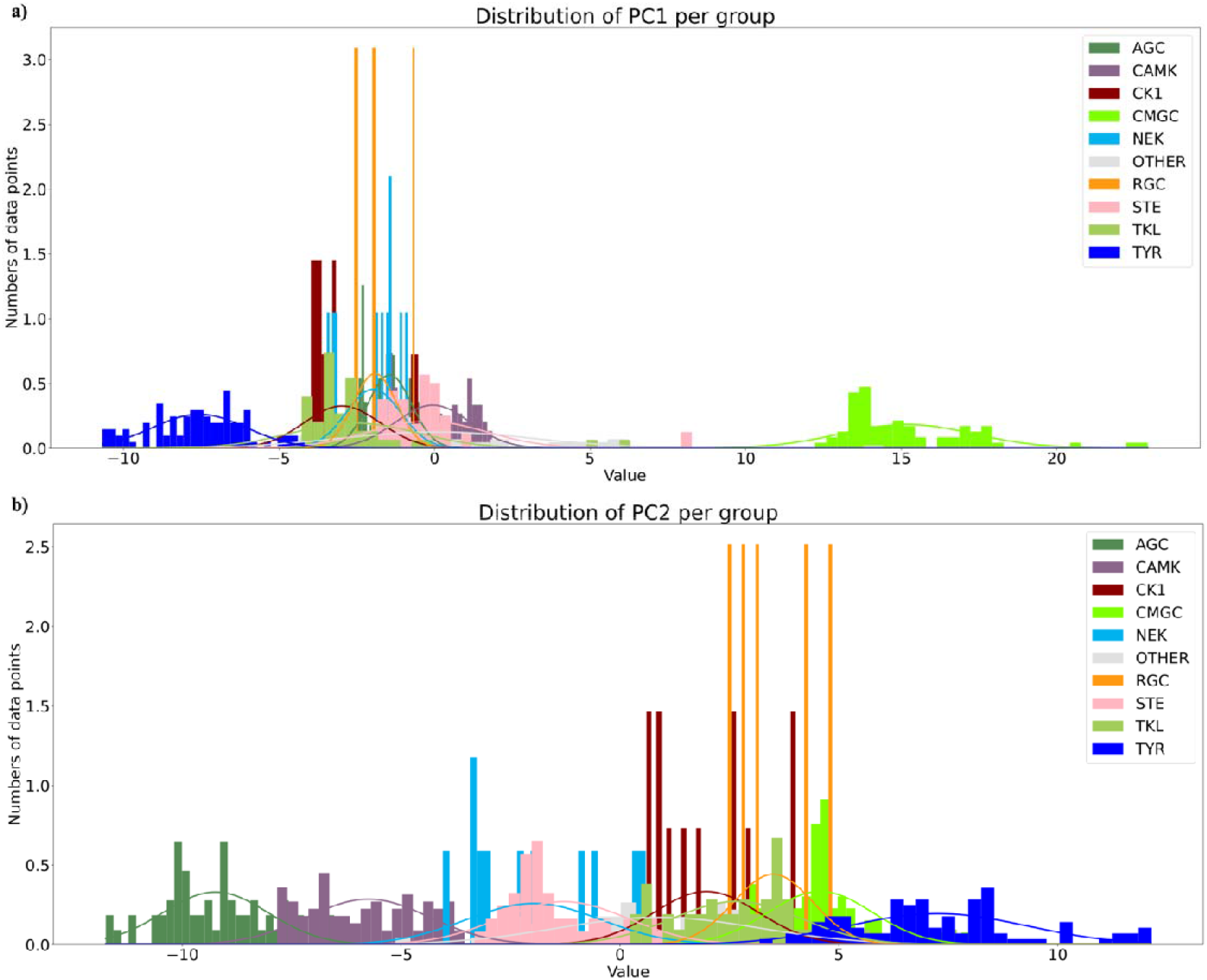
Two first PC distributions. **a)** PC1 and **b)** PC2 distributions, color coded based on KinCoRe classes. X-axis represents the PC values and the y-axis the number of data points, also known as the Probability Density.

A statistical analysis of the PCs, using a randomization test of PC weights multiplied by residue parameter values at each position, identified regions that separate specific classes of kinases. We did a pairwise randomization test of the distribution contribution of each position to PCs of each class versus the other classes, as well as each class versus all the other labeled kinases for the first 5 PCs. To complete the detection of significant position signals (p-value < 0.05), their effect size was measured by calculating Cohen’s d (d > 2.5) (**SI Table 4**). We performed an exploratory analysis of some positions with the highest effect sizes across all classes to find the environments that are specific to certain classes of kinases.

### Detection of key residues in “CMGC” proteins

Position 1912 for “CMGC” versus all the other labeled kinases, has a d > 20 for an AAindex charge parameter (**SI Figure 4**). It is exclusively occupied by arginine or lysine in the “CMGC” class, suggesting the critical role of the positive charge. In contrast, this position is generally occupied by uncharged or non-polar residues in other classes, with a few exceptions: LRRK2 from “TKL”, where position 1912 is arginine, and LRRK1 from “TKL” and STK11 from “CAMK” where it is occupied by glutamine.

As protein kinase structural domains are defined by molecular interactions between their residues, it is crucial to examine the surrounding environment when studying a specific position. For position 1912, we studied all residues within 5 radius, to ensure that all potential interactions, polar and non-polar, are captured. Position 1912 is in the activation loop region. To illustrate how “CMGC” is different in this region from the other classes, we selected two representative kinases, which exhibit “typical” residues around position 1912, and the corresponding experimental 3D structures (**Figure 5**): PDB ID 4MYG [38] for MAPK13 from “CMGC” (**Figure 5.a**); and PDB ID 6HHJ [41] for AKT1 from “AGC” (**Figure 5.b**). Of note, as the region is near a phosphorylation site for the “CMGC” class, we selected the 4MYG experimental structure to represent it since the corresponding threonine at position 1903 is phosphorylated (**Figure 5.a**).

**Figure 5.**
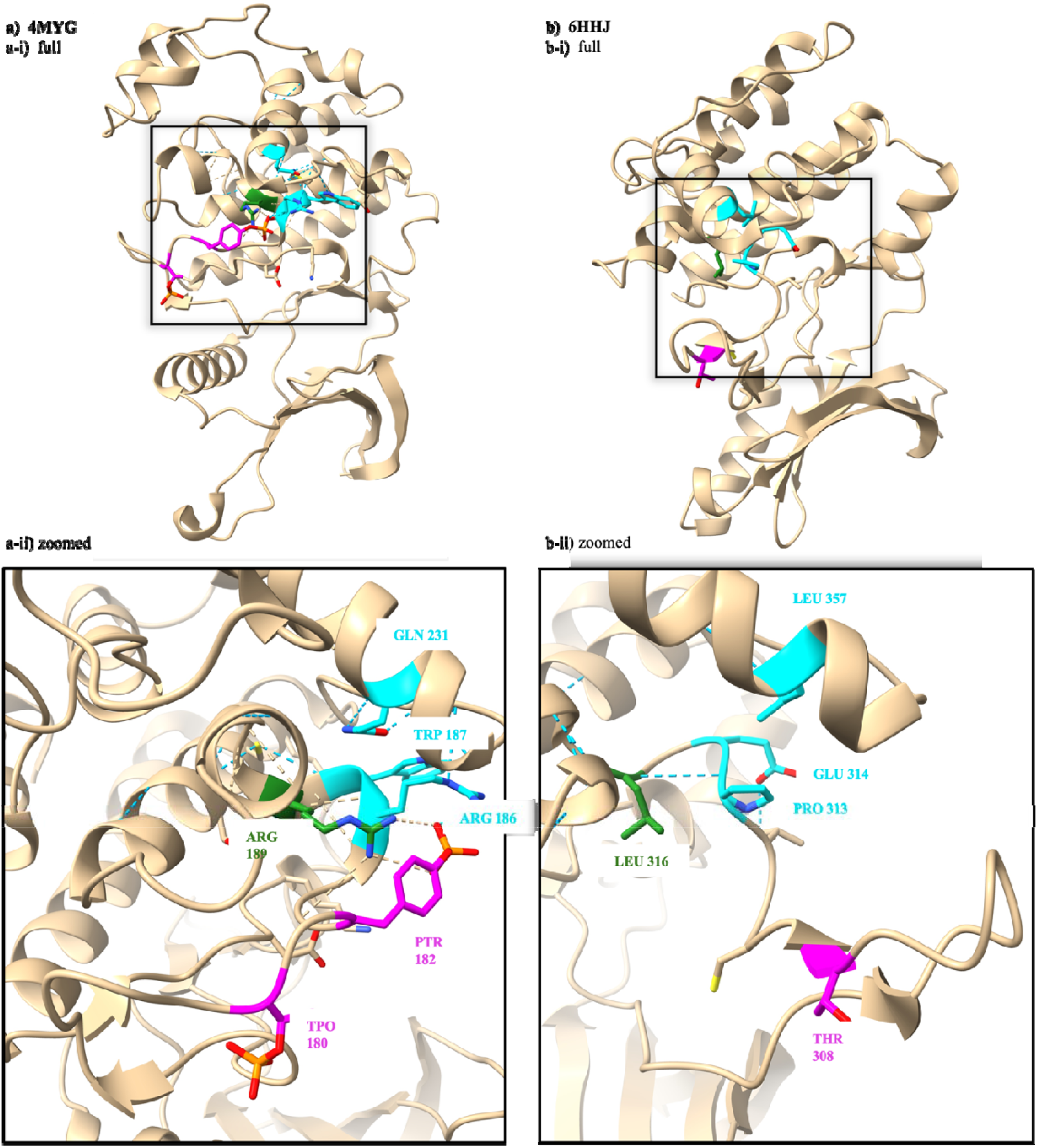
Structural analysis of position 1912 in kinase domains. **a)** Structure of MAPK13 labeled as “CMGC” (4MYG[38]); **b)** Structure of AKT1 labeled as “AGC” (6HHJ[41]; For both, **i)** is the full structure of the kinase domain and **ii)** is the zoomed in illustration of position 1912. The numbering corresponds to the protein canonical sequence: Position 1912 in alignment corresponds to different position numbers in each structure. In the structures we colored the position corresponding to alignment position 1912 in dark green. The positions colored in magenta are the phosphorylation sites close to this region based on the information acquired from PTM information in UniProt. Cyan residues correspond to the surrounding ones (distance < 5□) that have parameters with effect size higher than 1.5 and p-value < 0.05 in the randomization test: 1909, 1910, and 2053.

The KinCoRe alignment position numbers of important residues, displayed in **Figure 5**, are mapped with the standard protein residue numbers as provided by UniProt in **Table 1**. Residues surrounding position 1912 are 1909, 1910, and 2053. Position 1909 has significantly different size, charge, hydrophobicity, and polarity in the “CMGC” class compared to the corresponding one in the other classes (**SI Figure 5**). Position 1910 has different hydration and hydrophobicity (**SI Figure 6**). Position 2053 is different in terms of hydrophobicity, hydrophilicity, and polarity (**SI Figure 7**).

**Figure 6.**
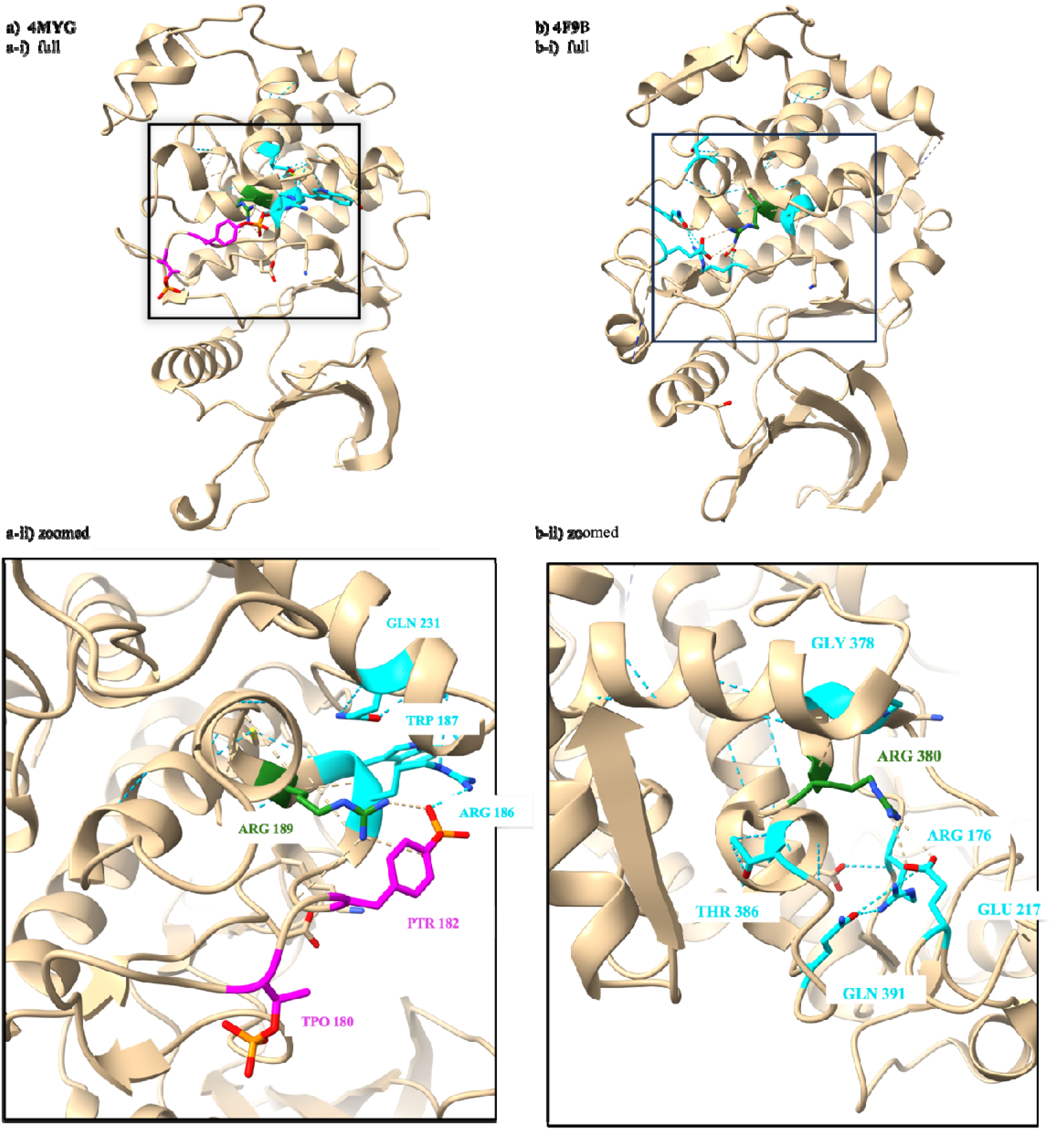
Side by side representation of structures of a CMGC kinase and an unlabeled kinase placed in the CMGC group. **a)** structure of phosphorylated kinase domain of MAPK13 labeled as “CMGC” (**4MYG) b) 4F9B**[42] **of unlabeled kinase CDC7 unanimously predicted in the CMGC class by all models.** Alignment position 1912 is shown in dark green. Residues colored in cyan have effect sizes > 1.5 for “CMGC” versus all the other labeled kinases. All non-carbon atoms are colored by atom type. The blue dashed line shows hydrogen bonds between atoms in this region. **i)** the whole kinase domain and **ii)** zoomed in on the region of interest, position 1912. The corresponding residues are labeled accordingly

**Table 1.**
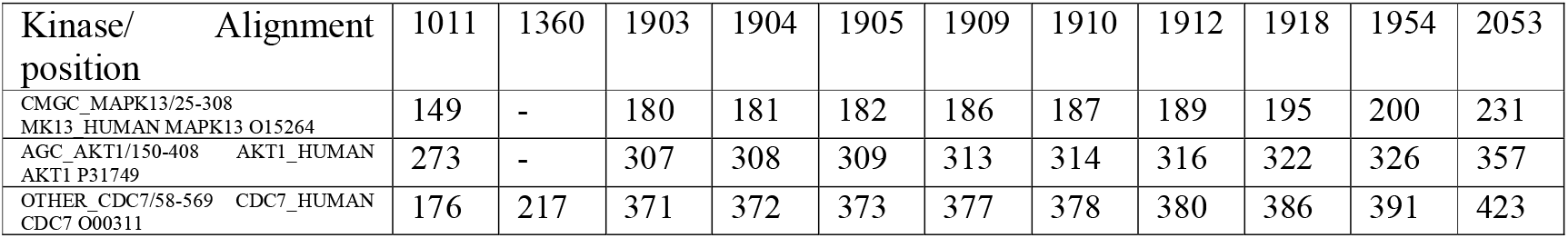
KinCoRe alignment positions mapped to the corresponding residue numbers in the mentioned kinases. The magenta color corresponds to phosphorylation sites. Cyan is for positions within the 1912 position that have a high effect size.

**Figure 5.a** displays a phosphorylated “CMGC” kinase domain of MAPK13. The phosphorylation of both Thr1903 and Tyr1905, colored in magenta, leads to kinase activation. **Figure 5.b** shows the AGC kinase domain of AKT1. Comparing the two structures, we see that Arg1912 makes hydrogen bonds with the phosphorylated Tyr1905 in MAPK13. A lot of kinases that belong to “CMGC” need to be phosphorylated at positions 1903 and 1905 to be activated^8^, which is not the case in this region for other classes. Available data indicates that the physico-chemical properties of the residues at position 1912 and its 3D environment play a role in stabilizing position 1905 phosphorylation, as shown by polar interactions between Arg1912 and phosphorylated Tyr1905 in the MAPK13 structure (**Figure 5.a**). These “CMGC”s have the typical “CMGC” structure in this region as well. “CMGC”s that do not have the typical “CMGC” residues are only phosphorylated at position 1903, and at position 1904 in other kinase classes. It should be noted that for the “CMGC” kinase, GSK3A, which is not dually phosphorylated, also carries a phosphorylation site at position 1905.

### Classification predictions of the unclassified kinases and structure-based validation

There are 66 unclassified kinases, labeled “OTHER” in KinCoRe. An ensemble model of supervised classifications using logistic regression, random forest, and Gaussian Naïve Bayes model was trained to map the PCA transformed kinase representations with class labels to their corresponding classes. We also used our unsupervised union k-means clustering to map kinases, where a data point is labeled as the highest number of kinase classes occupying a cluster (see **SI Table 5** for all the predictions along with the confidence score results). It is important to note that the models predict the class based on the highest probability of the corresponding class. Since no class is defined as NULL, a prediction is made for all data points. However, further investigation of the predictions should be made, by considering both the confidence score and the class-defining regions identified earlier through the randomization test.

There are 4 kinases from the “OTHER” class that are classified unanimously across all the models, with a confidence score of 7: CDC7, which is placed in “CMGC”. In the unsupervised model, the cluster assignment is 3, which is one of the clusters with the highest confidence. ULK1, ULK2, and ULK3 are grouped into “CAMK” (**SI Table 5**). As shown before, position 1912 and its surrounding environment constitute a distinct region among the kinase groups, which is very specific to “CMGC”. This position is occupied by arginine in 95% of “CMGC” kinases and only in “CMGC”. In CDC7, this position is occupied by arginine. To investigate the coherence of this classification, functional and structural analysis were performed (**Figure 6**).

The structure of AKT1 (**Figure 5.b**) shows a “typical” kinase, i.e., with residues more frequently present in this region across all the other labeled kinases. The KinCoRe alignment position numbers of important residues, displayed in *Error! Reference source not found.*, are mapped with the standard protein residue numbers as provided by UniProt in **Table 1** for MAPK13, AKT1 and CDC7, representing the “CMGC”, “AGC” and “OTHER” classes, respectively. In CDC7, position 1912 is very similar to the CMGC, which differs from kinases not labeled as “CMGC”, where this position is more exposed to the solvent (**Figure *6*)**. In “CMGC”, position 1912 is stabilized by the phosphorylated Tyr1905, and Arg1909 (*Error! Reference source not found.***.a**). In CDC7, this stabilization is done by Gln1954, Glu1360, and Arg1011 (**Figure 6.b**). These stabilization effects are not observed in other kinases, represented by AKT1 (**Figure 5.b**). We have shown more significant positions for CMGC in SI Figure 10, along with a side by side comparison of CDC7, a CMGC, and an AGC kinase to highlight the structural similarity of CDC7 and CMGC.

## Discussion

Using KinCoRe MSA and AAindex parameter residue properties, we studied kinases and their classes in a supervised and unsupervised manner. Most proteins have multiple domains. In our study, the range of the sequence that exists in the kinase domain of the protein is only chosen and analyzed. Each domain contributes to the biochemical role in a certain way. In this study, we focus exclusively on the sequence region corresponding to the kinase domain. While proteins sharing a highly similar kinase domain may perform very different biological functions at the organismal or cellular level, these differences often arise from the presence of other domains. By restricting the analysis to the kinase domain, our domain-centric approach provides a more fine-grained characterization. In the unsupervised method, we provided classifications that was only based on properties of position occupants. To determine the optimal number of clusters, we analyzed the PCA-transformed kinase representations with *k*-values ranging from 2 to 30, calculating silhouette scores, and detected local maxima at *k*= 11, 14, and 18. To assess the consistency of clusters across different *ks*, pairwise confusion matrices were generated, which highlighted the relationships between clusters at different *ks*, identifying coherent and conflicted clusters. An overall clustering was then created by combining the data points from the *ks*, resulting in 24 clusters with varying levels of consistency. The final analysis demonstrated that 363 kinases were assigned to consistent clusters, while 134 were placed in inconsistent clusters. The union of all *ks* clusters was compared to previously assigned class labels, revealing that 90% of the data points overlapped with known kinase classes. This unsupervised classification was further validated by comparing intra- and inter-cluster similarities, which showed that union clustering captured substantial group differences compared to the KinCoRe MSA, leading to a more precise classification. We used this unsupervised classification to propose a classification for previously unclassified kinases, labeled “OTHER” in KinCoRe (**SI Table 4**). Obviously, such predictions would have to be validated experimentally.

In the supervised portion of the study, we identified positions that contribute to statistically significant property differences between each kinase class using a permutation-based randomization test and effect size analysis. The structural analysis of these positions and their environment provides a better understanding of the role of these regions, which have been shown to be specific to classes of kinases. We have shown that some of these regions directly influence the kinase activity, like position 1912 in “CMGC”, which stabilizes the phosphorylation site at position 1905. At the physico-chemical level within the kinase domain, CDC7 shares greater similarity with “CMGC” kinases than with other kinase groups, as well as ULK1 and ULK3 to “CAMK”. Our framework captures global patterns rather than lineage-defining motifs. So, it reveals similarities that are orthogonal to traditional classification schemes. Our results do not redefine kinase group membership but instead map kinases into a physico-chemical similarity space, within which CDC7 exhibits a CMGC-like constraint profile despite its distinct evolutionary origin. The CMGC-arginine position is shown since it is statically significant to “CMGC”. However, our multi-variant classification method relies on all the positions to place each datapoint to a class and not only on position 1912. Although PTM information is currently incomplete for all “CMGC” kinases, our approach has allowed us to identify the uniqueness of this region, which was later confirmed through structural analysis. This highlights the strength of our approach in identifying key regions specific to certain kinase classes, which can be applied, for example, to the study of PTMs, such as phosphorylation sites.

To conclude, we developed an ensemble model that used the PCA-transformed KinCoRe alignment AAindex descriptors to predict kinase main classes. The model showed high performance in 10-fold cross-validation classification. We then applied this model to classify unclassed kinases, labeled as “OTHER” and further validated them using the class-specific positions that were found in the randomization test.

This study introduces a new approach for defining kinase classes and detecting their key differences. Although kinases perform the same enzymatic reaction, their sequences and 3D structure influence their functional mechanisms. While these proteins are generally well-studied, many still lack detailed information, particularly regarding PTMs. The identification of class-specific regions can help unravel their functional significance. In this article, we described one position with a key functional impact for “CMGC”. However, a substantial list of other key positions could be found, across all classes, which are presented in this article (**SI Table 4**). All this new information will contribute to a better understanding of kinase function in a more specific way, which can be useful in the developing new and more specific ligands and the discovering of new potential targets.

To demonstrate the universality of our method, we applied the same procedure to the small GTPase group. **SI Figure 11** shows clusters of the PCA-transformed representations. **SI Figure 12** demonstrates visually the difference in PC1 and PC2 values for different classes. Beyond improving kinase classification, our results demonstrate that this approach holds great potential to enhance our understanding of not only kinases but also other proteins and their domains, by providing deeper insights into key regions and their functional roles. These insights acquired can guide the design of ligands tailored to specific domain classes, in this case kinases, and help study mutations that may influence domain activity or substrate binding, ultimately advancing both therapeutic and functional research.

## Methods

All scripts were written in Python 3.9.7 using the following libraries: NumPy 1.20.3 [43], Pickle HIGHEST_PROTOCOL5, Matplotlib 3.4.3 [44], Seaborn 0.11.2 [45], Scikit-learn 0.24.2 [46] and SciPy 1.7.1 [47]. UMAP 0.5.3 [39] was used for visualizing the different kinase representations.

UCSF ChimeraX 1.7 [48] was used to perform the structural analysis and to generate images.

### Data extraction

The KinCoRe structure-based MSA of 497 kinases is composed of 2218 amino-acid positions [23]. The sequences are annotated with gene names, corresponding UniProt IDs, residues present in the kinase domain, and main classes in the protein kinase classifications.

The experimental 3D structures of kinases were extracted from the Protein Data Bank (PDB) [49]. For kinases with multiple structures, the one with the highest coverage and highest resolution has been chosen, with the highest coverage taking priority. Only experimental 3D structures with no mutation in the kinase domain were kept, leading to 227 structures.

### Kinase representation

The AAindex database [40] provides numerical indices for various physico-chemical and biochemical properties of amino acids. The indices represent amino acid size, charge, hydration, hydrophilicity, hydrophobicity, and polarity. We used 30 property descriptors for each amino acid in the MSA, after normalizing their values [40]. We added one descriptor as gap indicator. Each kinase is represented by a matrix of 2218 columns, each corresponding to an amino-acid position in the MSA, and 31 rows corresponding to AAindex property descriptors and alignment gap. As a kinase domain generally consists of an average of 275 residues, most of the 2218 positions for a given kinase in the MSA contain gaps and their AAindex values were set to zero.

### Principal component analysis (PCA) and randomization test

PCA was performed on kinase representations using PCA implementation from Scikit-learn [46]. There are crucial regions with specific properties that define the function of a kinase, and consequently which class it belongs to. To find regions and property parameters contributing to different principal components (PC) ranges for different classes, we calculated the dot product of PC weights to the original kinase representation for all kinases. These values were compared to each other for kinase classes by randomization tests to find regions with significantly different contributions.

Randomization test is a nonparametric approach where the data is randomly shuffled between two samples many times to create a null distribution. The observed statistic (median used here) is then compared to the null distribution to assess if the two samples come from the same distribution. The p-value indicates the probability of observing the test statistic under the null hypothesis. A threshold of p-value < 0.05 was chosen to reject the null hypothesis— i.e., to find positions that display significant difference between classes. The randomization test was performed 1000 times, for all class pairs and for kinases within each class versus all the other kinases, excluding the kinases classified as “OTHER”. At each position, a list of the residue parameter values multiplied by its PCA weight was calculated for all kinases. Given two kinase classes, a randomization test was performed to determine if the values are most likely from the same distribution or not. The positions with significantly different values were then displayed and studied. These values were calculated for regions with occupancy higher than 0.5. The occupancy was defined as the proportion of kinases that have a residue at a specific position. Importantly, a p-value on its own is not enough to determine if two classes are significantly different. The effect size must be considered as well, since with large enough sample sizes, it is highly likely that very small p-values are obtained [50]. Additionally, it is important to correct for the increased risk of false positives [51] when multiple hypothesis tests are performed simultaneously. We used the correction of the Bonferroni method [52]. To quantify the effect size between two classes, we calculated the Cohen’s *d* value as:

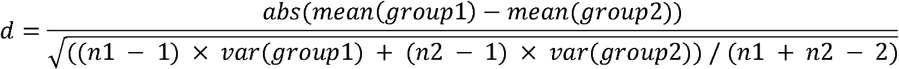

Where n1 = length of class1 and n2 = length of class2. Usually, Cohen’s d is interpreted as follows: d < 0.2 corresponds to a small effect size, d ∼ 0.5 to a medium effect size, and d > 0.8 to a large effect size, and d > 1.5 to a very strong effect [53,54]. However, it is also mentioned that these are the rule of thumb values and usually the value thresholds are determined based on the context. We analyzed the positions that have occupancy > 0.5, p-value < 0.05, and Cohen’s d > 2.5 on PC1 up to PC5 to unravel positions that are significantly different in AAindex parameters in a kinase family compared to the other families, which we will name class identifiers.

We applied the same method to another superfamily to test for the universality of this procedure. We chose small GTPase family. There are 162 proteins assigned to this group. The corresponding human reviewed sequences were retreived from UniProt and aligned using Clustal Omega [55]. Although a more meticulous alignment can contribute to the accuracy of the study, the available alignment techniques can also be used in this pipeline. We repeated the same steps to show the clusters of this family, when represented with AAindex descriptors (**SI Figure 11**), as well as different distributions of PC1 and PC2 values (**SI Figure 12**).

### Unsupervised *k*-means clustering

The PCA-transformed kinase representations are used as input for unsupervised clustering using *k*-means. The best number of *k* is determined by running the clustering algorithm for *k* from 2 to 30. The silhouette score for each different *k* cluster assignments are calculated. The best *k*s are the local maxima in this plot (**Figure 3.a-i**) i.e., *k*=11, 14, and 18.

We got the cluster assignments for all three different *k*s. The consistency between different *k* values were inferred as data points that belong to the same cluster for different *k*s. Of course for higher *k*s we accept if data points from a cluster of a lower *k* are divided into sub-clusters. We consider the cluster inconsistent if data points are clustered together for a higher *k*, but are divided for a lower *k*. Since we observed some inconsistencies between different *k*s, we presented a union clustering based on the combination of the cluster assignments for the three *k*s. There are 24 unique combinations across different *k*s.

To measure our union cluster confidence, we came up with a score. Each cluster is assigned a score of 10 in the beginning. The confidence score remains 10 if the cluster contains the same data points for *k=*11, 14, and 18. If for higher *k*s, we see a cluster of the same data points, but they are not in the same cluster for a lower *k*, then we will penalize the cluster score. The amount of penalizing is: -2 if consistent for *k=*11 and 18 but not 14; -3 if consistent for *k*=14 and 18 but not 11; and -5 if inconsistent between *k*=11 and 14. In the end, the union clusters have a score ranging from 5 to 10 (**SI Table 1**). We achnowledge that the union cluster scores are a measure of the quality of the union clusters based on the coherence between different *k*s and should not be presented as an absolute measure of the cluster quality.

### Defining residue environment for the structural analysis

The 227 experimental 3D structures of kinase domains were used to identify and study the environment of a specific residue at 5Å radius. This radius accounts for interactions such as hydrogen bonds, van der Waals forces, and salt bridges, which are crucial for maintaining protein stability and its activity, and this threshold is widely used in various studies [56,57]. A residue is defined as being in the vicinity of a reference residue, and included in the analysis, if at least one of its heavy atoms had a distance lower or equal than 5A to any atom of the reference residue.

### Logistic regression to map unclassified kinases

Given that we have unlabeled data, i.e., the class “OTHER”, it is of interest to study the probability array of kinases that belong to this class. To map unclassified kinases to a main class, Logistic Regression (LR), Random Forest (RF) and Gaussian Naïve Bayes (GNB) [58,59] models with various hyperparameters (**SI Table 6**) optimized via GridSearchCV from scikit-learn. The model performance was evaluated based on the highest average Matthews correlation coefficient (MCC) from 10-fold cross validation.

To avoid overfitting, it is important to determine the optimal number of PCs to include. The grid search was performed considering 1 to 20 PCs, and the average MCC of the best-performing hyperparameters for each model was calculated (0). Model performance began to plateau from 5 PCs onward, which served as the basis for training the final models. The hyperparameters chosen are C = 1 for LR, n_estimators = 300 and max_depth = 30 for RF, var_smoothing = 1e-9 for GNB (in bold in the **SI Table 6**). The mean MCC and accuracy across the folds are shown in **SI Figure 8**. The performance of the models in detail is shown in **SI Table 7**.

We applied the three models (LR, RF and GNB) to predict the class of the kinases labeled “OTHER” in a consensus way. Unanimous class predictions were analyzed and explored if they are valid in view of the class specific randomization test results (**SI Table 5**).

## Supporting information

Supplementary Information - Revised

## Acknowledgement

The authors would like to thank the University of Lausanne, the Ludwig Institute for Cancer Research Lausanne and the foundation Donase.

## Key points

- This work highlights the importance of physico-chemical residue properties in describing protein sequences. Its application to kinase members has demonstrated improved classification performance, offering greater precision compared to traditional phylogenetic approaches.
- By analyzing the PC distribution of physico-chemical properties, the method reveals class-specific sequence regions and positions, as evidenced by statistically significant differences between classes.
- This novel approach is applicable to all protein types and can be used to detect PTM sites or identify critical positions within protein sequences.

