## Supplementary Information - Revised for "Novel universal domain-centric method for protein classification"

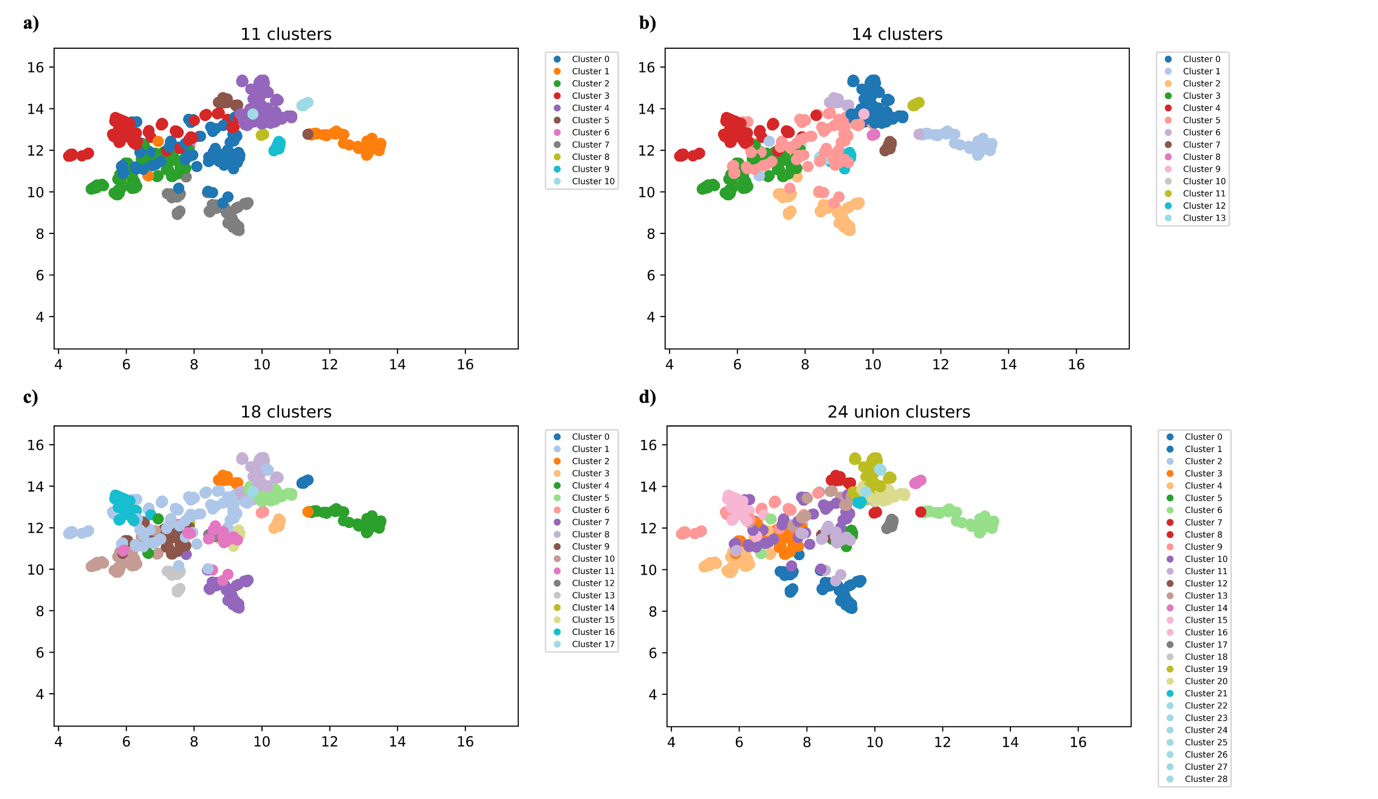

**SI Figure 1. UMAP visualization of PCA-transformed kinases color coded based on the k-means clustering. a)** k = 11, **b)** k = 14, **c)** k = 18, and d) k=24 union clusters

**SI Table 1. Union cluster assignments**. The new assignments are based on the assignment union of *k*=11, 15, and 17. We have 26 unique combinations of these three unsupervised k-means clustering.

| New cluster  Assignment (k=24) | Confidence  score | *k*=11 | *k*=14 | *k*=18 |
| --- | --- | --- | --- | --- |
| 0 | 10 | 7 | 2 | 7 |
| 1 | 10 | 7 | 2 | 13 |
| 2 | 10 | 6 | 10 | 14 |
| 3 | 10 | 2 | 3 | 9 |
| 4 | 10 | 2 | 3 | 10 |
| 5 | 10 | 0 | 12 | 15 |
| 6 | 10 | 1 | 1 | 4 |
| 7 | 10 | 5 | 6 | 2 |
| 8 | 10 | 8 | 8 | 6 |
| 9 | 5 | 3 | 4 | 1 |
| 10 | 5 | 0 | 5 | 1 |
| 11 | 10 | 0 | 5 | 11 |
| 12 | 10 | 0 | 13 | 12 |
| 13 | 5 | 3 | 5 | 1 |
| 14 | 10 | 10 | 11 | 0 |
| 15 | 5 | 2 | 3 | 1 |
| 16 | 10 | 3 | 4 | 16 |
| 17 | 10 | 9 | 7 | 3 |
| 18 | 8 | 4 | 5 | 5 |
| 19 | 10 | 4 | 0 | 8 |
| 20 | 8 | 4 | 0 | 5 |
| 21 | 5 | 4 | 5 | 1 |
| 22 | 5 | 4 | 0 | 1 |
| 23 | 10 | 10 | 9 | 17 |

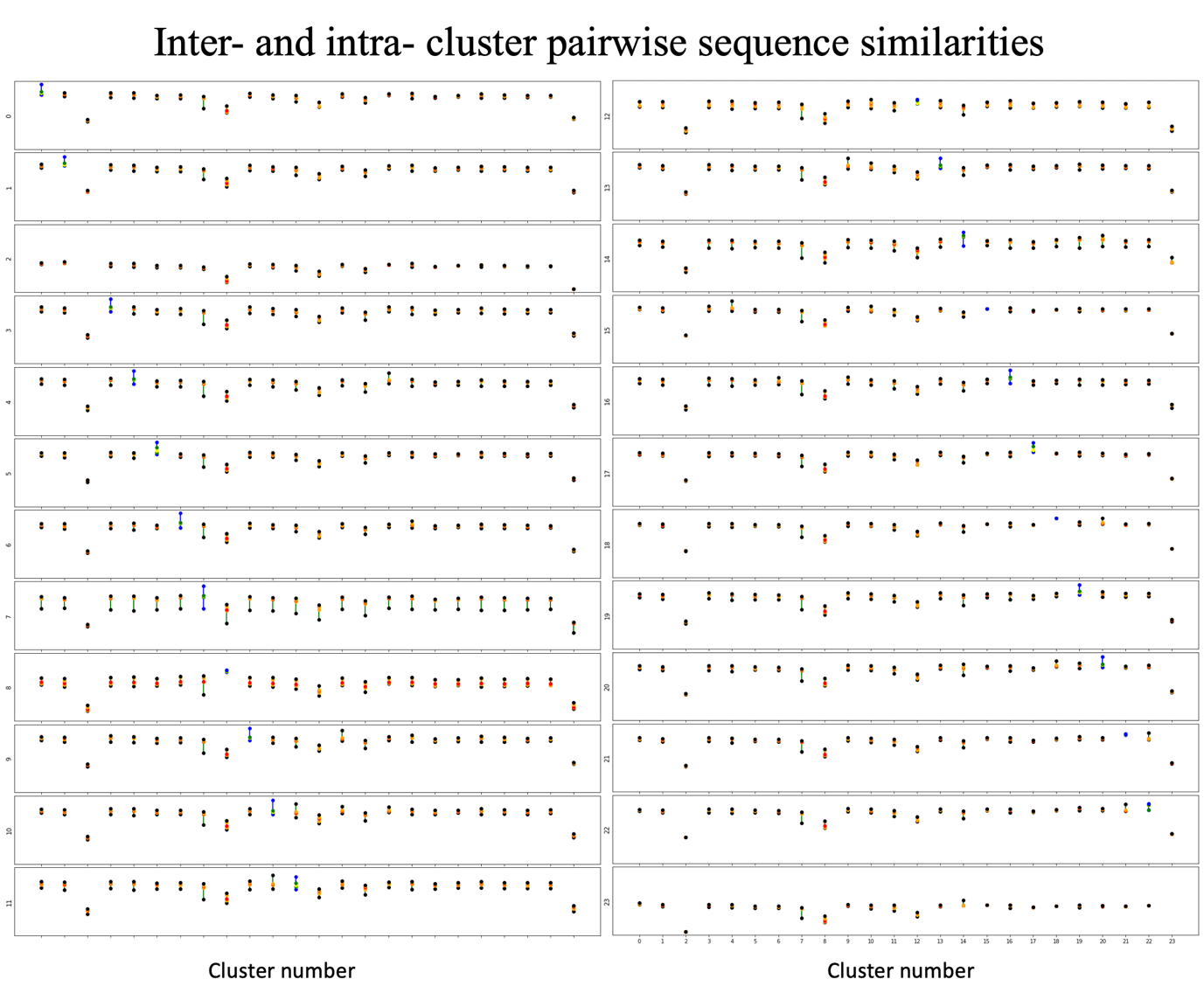

**SI Figure 2. Inter- and intra-cluster pairwise sequence similarity for the kinases within and between different clusters in our union k-means clustering**. Each sub-Fig. corresponds to a cluster highlighted in the y-axis. Each vertical line corresponds to the cluster number to which the data is compared to on the x-axis. Each vertical line is marked by 25 and 75 percentile and the median of the similarities between the two cluster data points. The black vertical lines are inter-cluster similarities and the blue ones are the intra-cluster similarities.

**SI Table 2. ANOVA test results for cluster difference.** This table shows the how different each cluster pairwise difference is compared to the other cluster pairwise differences.

| **Cluster** | **F-statistics** | **P-value** |
| --- | --- | --- |
| 0 | 7069.42 | 0 |
| 1 | 1630.30 | 0 |
| 2 | nan | nan |
| 3 | 4904.02 | 0 |
| 4 | 5878.21 | 0 |
| 5 | 1166.11 | 0 |
| 6 | 5717.82 | 0 |
| 7 | 224.23 | 0 |
| 8 | 38.67 | 4.2816e-263 |
| 9 | 3408.47 | 0 |
| 10 | 4454.85 | 0 |
| 11 | 876.99 | 0 |
| 12 | 82.75 | 0 |
| 13 | 1855.34 | 0 |
| 14 | 335.71 | 0 |
| 15 | 288.01 | 0 |
| 16 | 3499.20 | 0 |
| 17 | 1949.63 | 0 |
| 18 | 349.62 | 0 |
| 19 | 6370.72 | 0 |
| 20 | 4068.35 | 0 |
| 21 | 418.37 | 0 |
| 22 | 806.58 | 0 |
| 23 | nan | nan |

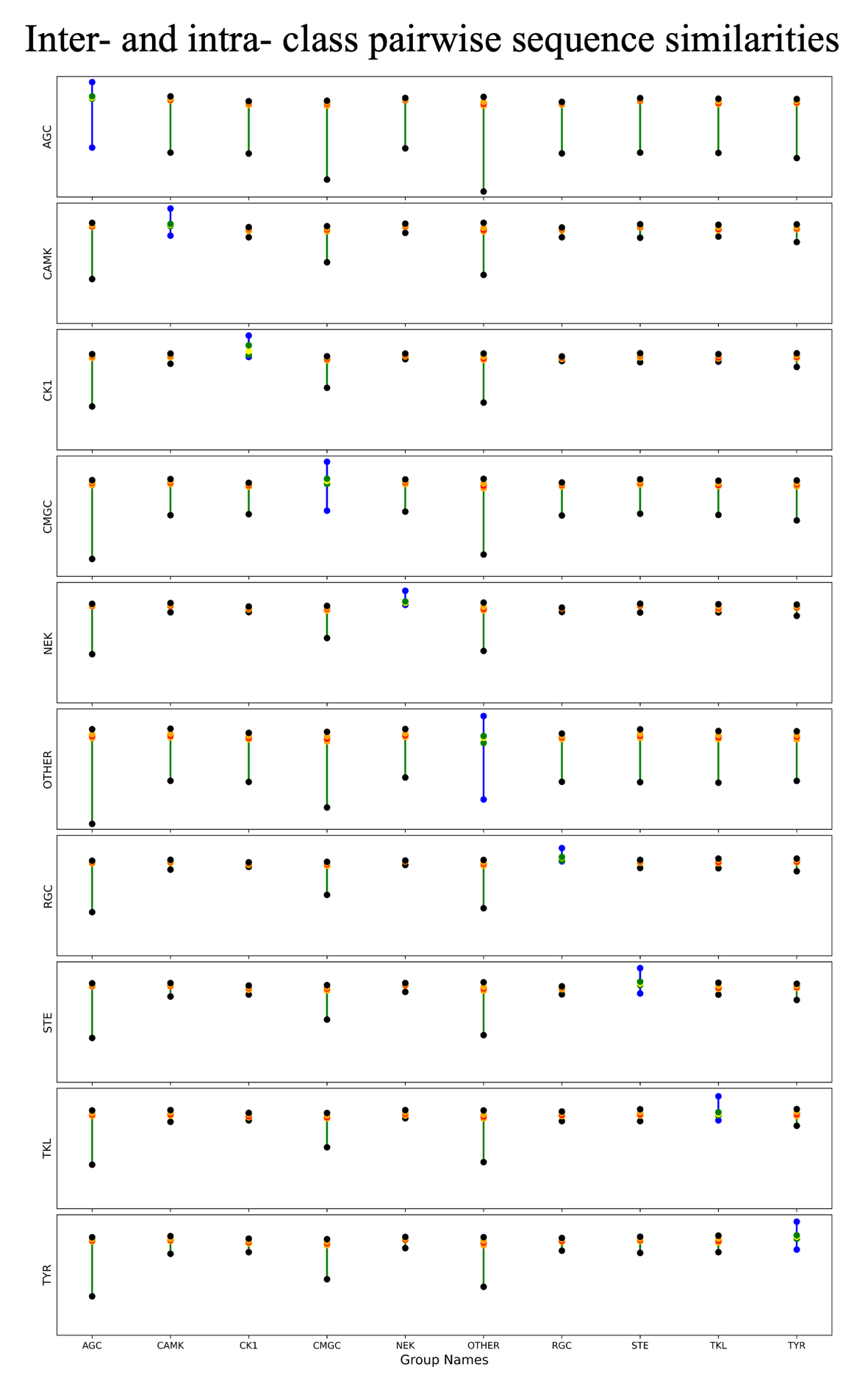

**SI Figure 3. Inter- and intra-cluster pairwise sequence similarity for the kinases within and between different classes in KinCoRe classifications**. Each sub-Fig. corresponds to a class highlighted in the y-axis. Each vertical line corresponds to the class to which the first class is compared to on the x-axis. Each vertical line is marked by 25 and 75 percentile and the median of the similarities between the two cluster data points. The black vertical lines are inter-cluster similarities and the blue ones are the intra-cluster similarities.

**SI Table 3. ANOVA test results for class difference.** ANOVA test result showing how different each class pairwise difference is compared to the other class pairwise differences.

| **Class** | **F-statistics** | **P-value** |
| --- | --- | --- |
| AGC | 324.85 | 0 |
| CAMK | 957.65 | 0 |
| CK1 | 119.19 | 0 |
| CMGC | 197.53 | 0 |
| NEK | 121.17 | 0 |
| OTHER | 106.57 | 0 |
| RGC | 46.19 | 9.55E-152 |
| STE | 458.67 | 0 |
| TKL | 343.22 | 0 |
| TYR | 1157.90 | 0 |

**SI Table 4.** The number of positions that are significantly dissimilar between kinases of different classes (randomization test corrected p-value < 0.05) with effect size Cohen’s d > 2.5.

| **Class** | **AGC** | **CAMK** | **CK1** | **CMGC** | **NEK** | **RGC** | **STE** | **TKL** | **TYR** | ***Vs.* all the other** |
| --- | --- | --- | --- | --- | --- | --- | --- | --- | --- | --- |
| **AGC** | - | 7 | 73 | 23 | 35 | 73 | 23 | 23 | 36 | 6 |
| **CAMK** | 7 | - | 51 | 11 | 19 | 48 | 11 | 15 | 25 | 1 |
| **CK1** | 72 | 50 | - | 64 | 11 | 4 | 51 | 44 | 77 | 31 |
| **CMGC** | 23 | 11 | 64 | - | 25 | 62 | 16 | 23 | 32 | 6 |
| **NEK** | 35 | 19 | 12 | 24 | - | 1 | 17 | 15 | 45 | 2 |
| **RGC** | 74 | 49 | 5 | 60 | 2 | - | 41 | 15 | 38 | 26 |
| **STE** | 23 | 11 | 52 | 17 | 18 | 42 | - | 6 | 29 | 2 |
| **TKL** | 23 | 15 | 45 | 23 | 15 | 13 | 6 | - | 8 | 1 |
| **TYR** | 36 | 25 | 77 | 32 | 45 | 40 | 30 | 8 | - | 9 |
| ***Vs.* all the other** | 6 | 1 | 31 | 6 | 2 | 26 | 2 | 1 | 9 | - |

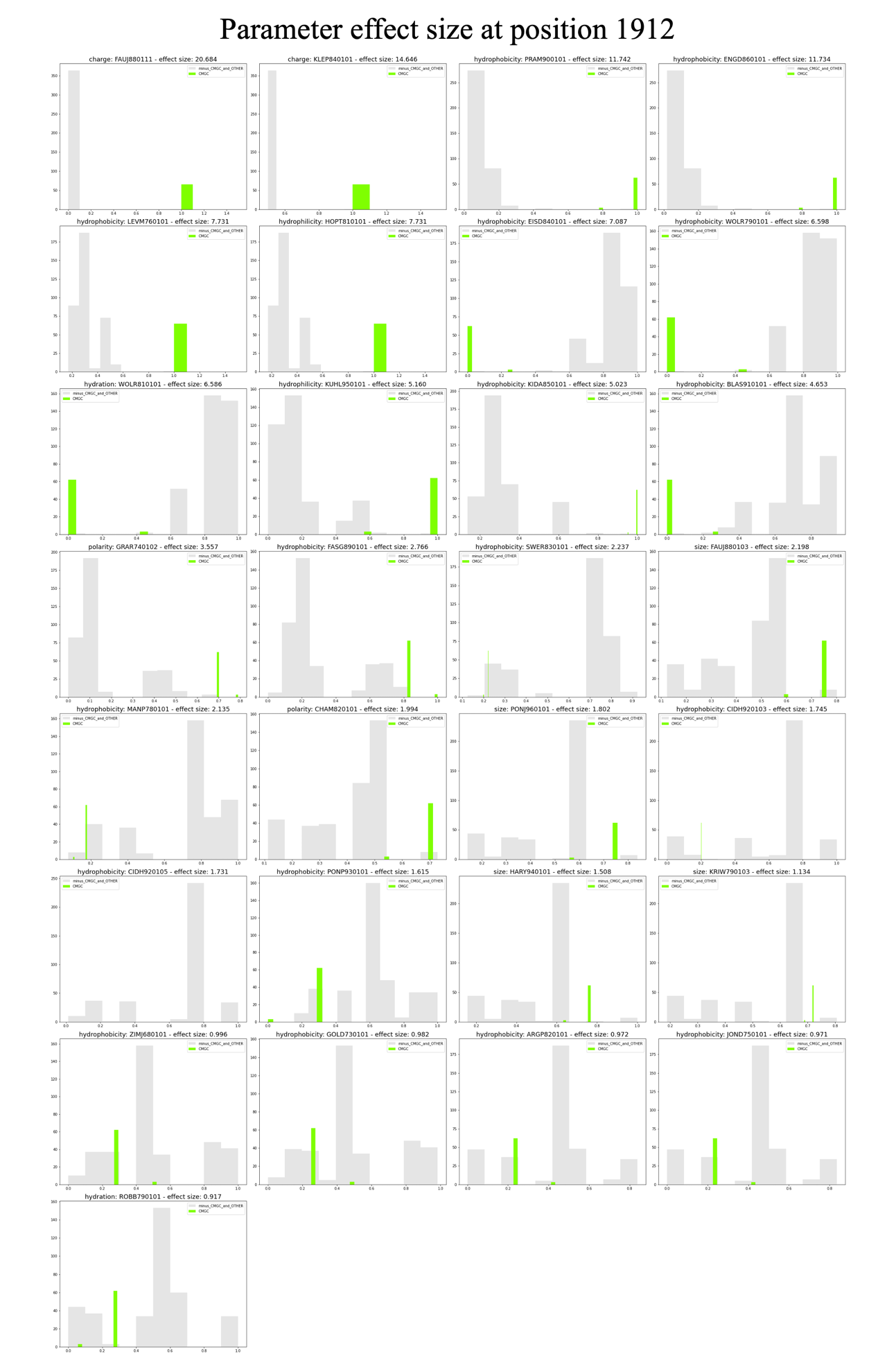

**SI Figure 4.** **Parameter effect size at position 1912**. Position 1912 is significantly different, with the highest effect size for CMGC kinases versus all the other kinases. These plots show different values corresponding to the residues at position 1912 in CMGC class (green) versus all the other labeled kinases (grey). As the effect size (shown in the title of the subplots) decreases, the separation between the two colors become less distinct.

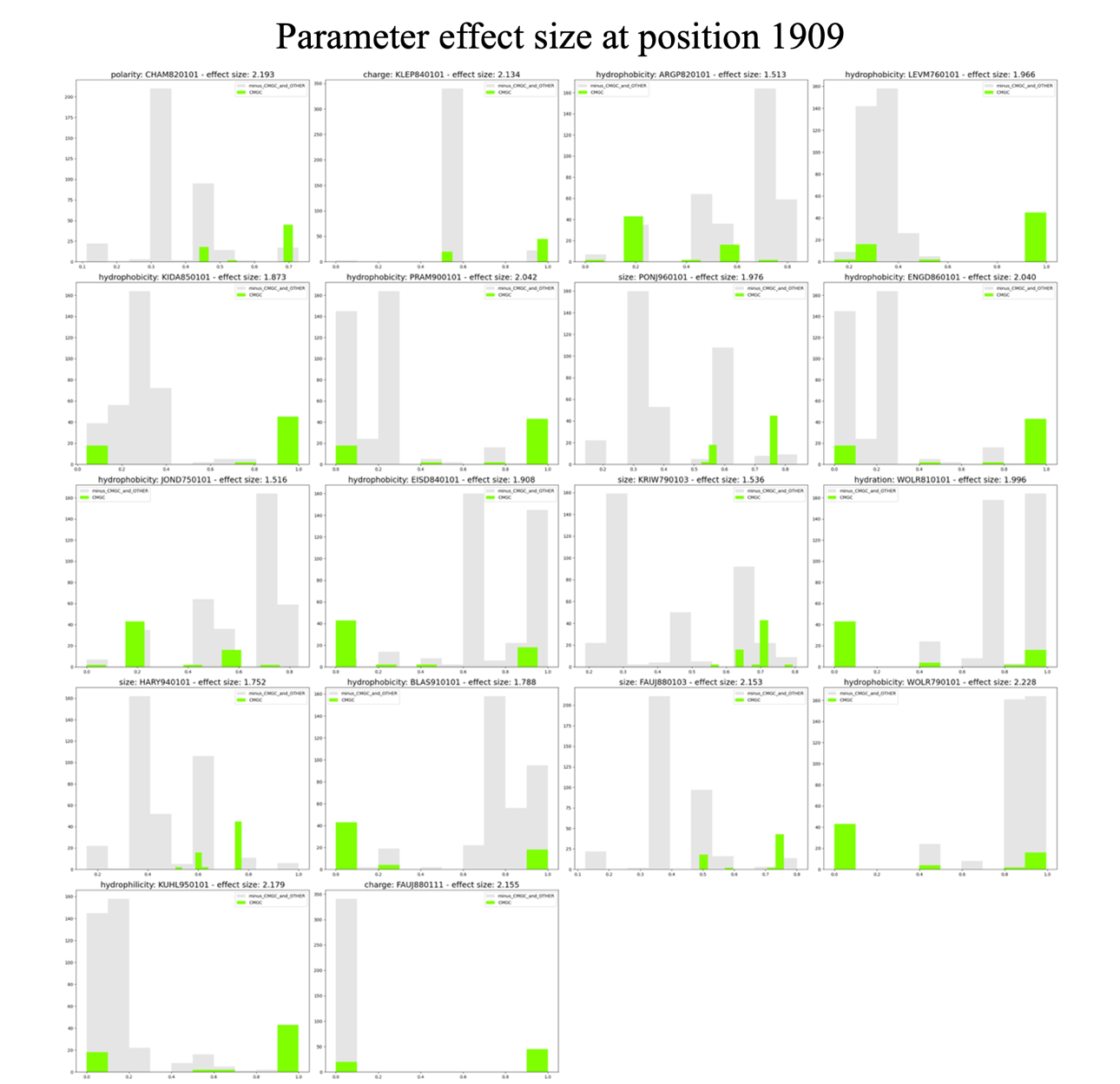

**SI Figure 5. Parameter effect size at position 1909**. Position 1909 is a significantly different position in the 5Å vicinity of position 1912 in CMGC versus all the other labeled kinases with AAindex parameters with effect size > 1.5. This Fig. shows the AAindex parameter values of residues occupying this position in CMGC kinases (in green) versus all the other labeled kinases (in grey). Position 1909 is mainly occupied by arginine (66%) followed by leucine (24%) in CMGC. Hydrophobicity parameters WOLR790101 and KUHL950101 have the highest effect size. Arginine in this position contributes to stabilizing a phosphorylated Y1905 leading to kinase activity. In the presence of a phosphorylation site in 1905, the presence of arginine at position 1909 is essential for stabilizing a phosphorylated kinase. This arginine is not needed in kinases where position 1905 is not a phosphorylation site, hence the leucine in the 24%.

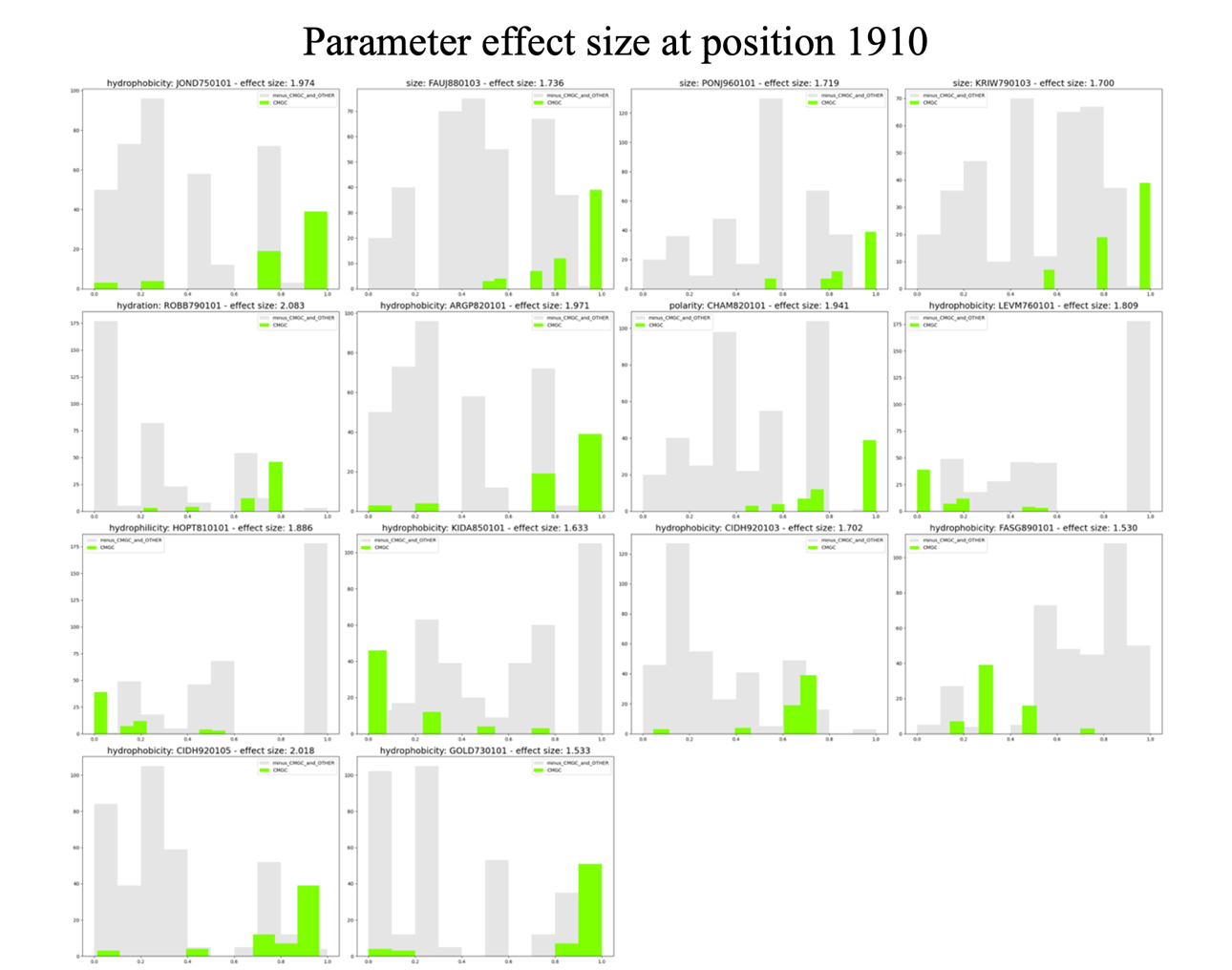

**SI Figure 6.** **Parameter effect size at position 1910**. AAindex parameter values of residues occupying position 1910 in CMGC kinases (in green) versus all the other labeled kinases (in grey). Position 1910, in the 5Å vicinity of position 1912, is a significantly different in CMGC versus all the other labeled kinases, with AAindex parameters with effect size > 1.5. Position 1910 is mainly occupied by tryptophan (60%), tyrosine (18%) and phenylalanine (10%) in CMGC. Hydration and hydrophobicity parameters ROBB790101 and CIDH920105 have the highest effect size. Glutamic acid, arginine, and lysine occupy most of the other labeled kinases at this position.

.
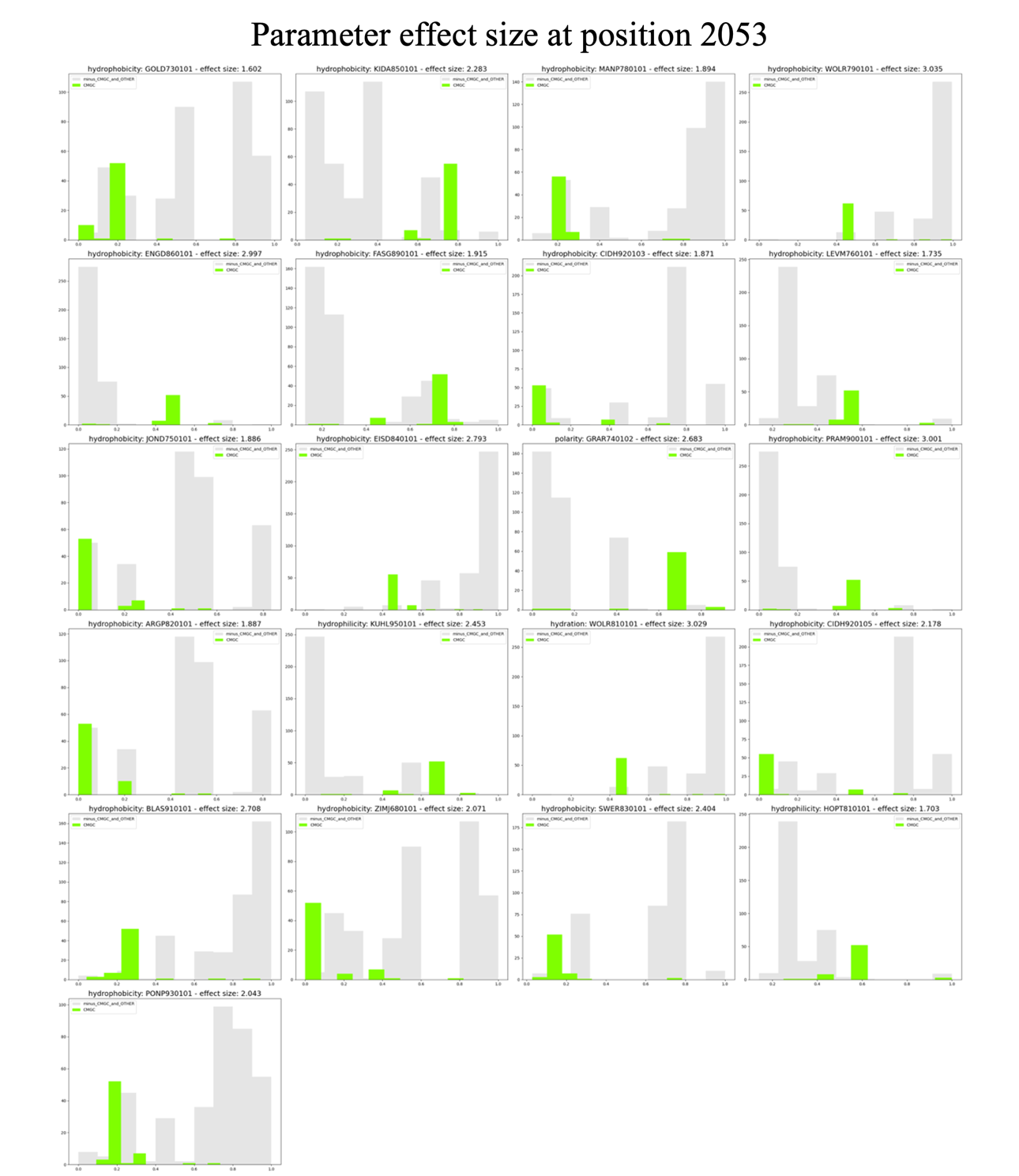

**SI Figure 7.** **Parameter effect size at position 2053**. AAindex parameter values of residues occupying position 2053 in CMGC kinases (in green) versus all the other labeled kinases (in grey). Position 2053, in the 5Å vicinity of position 2053, is a significantly different in CMGC versus all the other labeled kinases, with AAindex parameters with effect size > 1.5. Position 2053 is mainly occupied by glutamine (80%) and histidine (10%) in CMGC. The residues that occupy this position in all the other labeled kinases are primarily leucine, valine, isoleucine, etc. Hydrophobicity parameters have WOLR790101 and PRAM900101 as well as hydration parameters WOLR810101 have effect size > 3. Other parameters show high effect sizes as well, highlighting CMGC being very distinguished in this position versus all the other labeled kinases.

**SI Table 5. Class predictions for unclassified kinases and the confidence score of the predictions.** Supervised: Logistic Regression (LR), Random Forest (RF), Gaussian Naive Bayes (GNB). Unsupervised: Union k-means clustering.

| **Kinase** | **LR**  **prediction** | **LR**  **Score** | **RF**  **prediction** | RF Score | **GNB**  **prediction** | **GNB Score** | **Union**  **k-means**  **cluster** | **Union**  **k-means**  **cluster**  **assignment** |
| --- | --- | --- | --- | --- | --- | --- | --- | --- |
| **OTHER_AAK1/46-313 AAK1_HUMAN AAK1 Q2M2I8** | STE | 0.70 | STE | 0.52 | TKL | 1.00 | 10 | TKL |
| **OTHER_BMP2K/51-317 BMP2K_HUMAN BMP2K Q9NSY1** | STE | 0.67 | STE | 0.59 | TKL | 0.99 | 10 | TKL |
| **OTHER_BUB1/787-1056 BUB1_HUMAN BUB1 O43683** | CK1 | 1.00 | CK1 | 0.56 | CK1 | 0.99 | 11 | CAMK |
| **OTHER_BUB1B/766-1021 BUB1B_HUMAN BUB1B O60566** | CK1 | 1.00 | CK1 | 0.82 | CK1 | 1.00 | 11 | CAMK |
| **OTHER_CDC7/58-569 CDC7_HUMAN CDC7 O00311** | CMGC | 0.97 | CMGC | 0.84 | CMGC | 1.00 | 7 | CMGC |
| **OTHER_CHUK/15-309 IKKA_HUMAN CHUK O15111** | STE | 0.59 | NEK | 0.32 | TKL | 0.69 | 10 | TKL |
| **OTHER_DSTYK/648-908 DUSTY_HUMAN DSTYK Q6XUX3** | TKL | 0.91 | TKL | 0.67 | TKL | 0.61 | 10 | TKL |
| **OTHER_EIF2AK1/167-583 E2AK1_HUMAN EIF2AK1 Q9BQI3** | TKL | 0.98 | TKL | 0.87 | TKL | 1.00 | 12 | OTHER |
| **OTHER_EIF2AK2/267-538 E2AK2_HUMAN EIF2AK2 P19525** | TKL | 0.92 | TKL | 0.66 | RGC | 0.57 | 10 | TKL |
| **OTHER_EIF2AK3/593-1077 E2AK3_HUMAN EIF2AK3 Q9NZJ5** | TKL | 0.93 | TKL | 0.68 | TKL | 1.00 | 12 | OTHER |
| **OTHER_EIF2AK4_1/280-539 E2AK4_HUMAN EIF2AK4 Q9P2K8** | STE | 0.88 | STE | 0.57 | TKL | 1.00 | 11 | CAMK |
| **OTHER_EIF2AK4_2/590-1001 E2AK4_HUMAN EIF2AK4 Q9P2K8** | STE | 0.95 | STE | 0.60 | TKL | 1.00 | 12 | OTHER |
| **OTHER_ERN1/569-832 ERN1_HUMAN ERN1 O75460** | STE | 0.95 | STE | 0.56 | TKL | 0.98 | 10 | TKL |
| **OTHER_ERN2/518-781 ERN2_HUMAN ERN2 Q76MJ5** | STE | 0.97 | STE | 0.67 | TKL | 0.55 | 10 | TKL |
| **OTHER_GAK/40-315 GAK_HUMAN GAK O14976** | TKL | 0.80 | TKL | 0.65 | TKL | 0.61 | 10 | TKL |
| **OTHER_HASPIN/484-797 HASP_HUMAN GSG2 Q8TF76** | CK1 | 0.93 | CK1 | 0.67 | TKL | 1.00 | 11 | CAMK |
| **OTHER_IKBKB/15-308 IKKB_HUMAN IKBKB O14920** | STE | 0.57 | CMGC | 0.25 | TKL | 0.87 | 10 | TKL |
| **OTHER_IKBKE/9-304 IKKE_HUMAN IKBKE Q14164** | STE | 0.98 | STE | 0.42 | STE | 0.84 | 10 | TKL |
| **OTHER_MLKL/201-468 MLKL_HUMAN MLKL Q8NB16** | TKL | 0.90 | TKL | 0.85 | TKL | 1.00 | 10 | TKL |
| **OTHER_MOS/60-340 MOS_HUMAN MOS P00540** | TKL | 0.60 | RGC | 0.38 | TKL | 1.00 | 10 | TKL |
| **OTHER_NRBP1/64-327 NRBP_HUMAN NRBP1 Q9UHY1** | TKL | 0.54 | CK1 | 0.46 | CK1 | 0.97 | 10 | TKL |
| **OTHER_NRBP12/37-306 NRBP2_HUMAN NRBP2 Q9NSY0** | CK1 | 0.66 | CK1 | 0.68 | CK1 | 1.00 | 11 | CAMK |
| **OTHER_PAN3/480-751 PAN3_HUMAN PAN3 Q58A45** | CK1 | 0.92 | CK1 | 0.54 | CK1 | 0.93 | 11 | CAMK |
| **OTHER_PBK/32-320 TOPK_HUMAN PBK Q96KB5** | TKL | 0.87 | TKL | 0.77 | RGC | 0.75 | 10 | TKL |
| **OTHER_PDIK1L/8-331 PDK1L_HUMAN PDIK1L Q8N165** | STE | 0.92 | TKL | 0.67 | TKL | 1.00 | 11 | CAMK |
| **OTHER_PEAK1/1327-1665 PEAK1_HUMAN PEAK1 Q9H792** | CK1 | 1.00 | CK1 | 0.81 | CMGC | 0.93 | 11 | CAMK |
| **OTHER_PEAK3/171-397 PEAK3_HUMAN PEAK3 Q6ZS72** | CK1 | 1.00 | CK1 | 0.83 | CK1 | 1.00 | 11 | CAMK |
| **OTHER_PIK3R4/26-313 PI3R4_HUMAN PIK3R4 Q99570** | STE | 0.72 | STE | 0.57 | TKL | 1.00 | 10 | TKL |
| **OTHER_PINK1/156-509 PINK1_HUMAN PINK1 Q9BXM7** | TKL | 0.56 | CK1 | 0.68 | CK1 | 0.93 | 11 | CAMK |
| **OTHER_PKDCC/138-391 PKDCC_HUMAN PKDCC Q504Y2** | CK1 | 0.69 | CK1 | 0.68 | CK1 | 1.00 | 11 | CAMK |
| **OTHER_PKMYT1/110-359 PMYT1_HUMAN PKMYT1 Q99640** | STE | 0.55 | STE | 0.75 | STE | 0.94 | 13 | TKL |
| **OTHER_POMK/81-333 SG196_HUMAN POMK Q9H5K3** | TKL | 0.60 | TYR | 0.66 | TKL | 0.93 | 11 | CAMK |
| **OTHER_PRAG1/992-1327 PRAG1_HUMAN PRAG1 Q86YV5** | CK1 | 1.00 | CK1 | 0.80 | CMGC | 1.00 | 11 | CAMK |
| **OTHER_PXK/146-396 PXK_HUMAN PXK Q7Z7A4** | CK1 | 0.78 | CK1 | 0.48 | CK1 | 0.86 | 11 | CAMK |
| **OTHER_RNASEL/361-586 RN5A_HUMAN RNASEL Q05823** | CK1 | 0.57 | CK1 | 0.78 | CK1 | 0.97 | 11 | CAMK |
| **OTHER_RPS6KL1/150-539 RPKL1_HUMAN RPS6KL1 Q9Y6S9** | AGC | 1.00 | CMGC | 0.22 | CMGC | 1.00 | 14 | TYR |
| **OTHER_SBK1/53-315 SBK1_HUMAN SBK1 Q52WX2** | CAMK | 0.89 | STE | 0.48 | CAMK | 0.97 | 15 | OTHER |
| **OTHER_SBK2/62-327 SBK2_HUMAN SBK2 P0C263** | STE | 0.57 | STE | 0.55 | TKL | 0.99 | 10 | TKL |
| **OTHER_SBK3/43-306 SBK3_HUMAN SBK3 P0C264** | STE | 0.55 | STE | 0.57 | TKL | 0.96 | 10 | TKL |
| **OTHER_SCYL1/14-263 SCYL1_HUMAN SCYL1 Q96KG9** | STE | 0.72 | STE | 0.58 | TKL | 1.00 | 11 | CAMK |
| **OTHER_SCYL2/32-327 SCYL2_HUMAN SCYL2 Q6P3W7** | AGC | 0.88 | CAMK | 0.31 | TKL | 0.97 | 10 | TKL |
| **OTHER_SCYL3/11-245 PACE1_HUMAN SCYL3 Q8IZE3** | TKL | 0.81 | TKL | 0.55 | TKL | 0.82 | 11 | CAMK |
| **OTHER_STK16/20-292 STK16_HUMAN STK16 O75716** | TKL | 0.92 | TKL | 0.92 | TKL | 1.00 | 10 | TKL |
| **OTHER_STK31/710-972 STK31_HUMAN STK31 Q9BXU1** | AGC | 0.61 | STE | 0.47 | TKL | 1.00 | 11 | CAMK |
| **OTHER_STK35/202-529 STK35_HUMAN STK35 Q8TDR2** | STE | 0.95 | TKL | 0.72 | TKL | 1.00 | 11 | CAMK |
| **OTHER_STK36/4-254 STK36_HUMAN STK36 Q9NRP7** | CAMK | 0.49 | CAMK | 0.68 | NEK | 0.57 | 3 | CAMK |
| **OTHER_STKLD1/28-297 STKL1_HUMAN STKLD1 Q8NE28** | TKL | 0.71 | CK1 | 0.42 | TKL | 1.00 | 11 | CAMK |
| **OTHER_TBCK/1-273 TBCK_HUMAN TBCK Q8TEA7** | STE | 0.94 | TKL | 0.52 | STE | 0.88 | 13 | TKL |
| **OTHER_TBK1/7-304 TBK1_HUMAN TBK1 Q9UHD2** | STE | 0.99 | TKL | 0.62 | STE | 0.60 | 10 | TKL |
| **OTHER_TEX14/227-512 TEX14_HUMAN TEX14 Q8IWB6** | TYR | 0.95 | TYR | 0.48 | TYR | 0.74 | 10 | TKL |
| **OTHER_TLK1/456-734 TLK1_HUMAN TLK1 Q9UKI8** | STE | 0.94 | STE | 0.69 | STE | 0.94 | 10 | TKL |
| **OTHER_TLK2/462-741 TLK2_HUMAN TLK2 Q86UE8** | STE | 0.94 | STE | 0.69 | STE | 0.93 | 10 | TKL |
| **OTHER_TP53RK/33-253 PRPK_HUMAN TP53RK Q96S44** | TKL | 0.61 | CK1 | 0.61 | CK1 | 0.75 | 11 | CAMK |
| **OTHER_TTK/525-791 TTK_HUMAN TTK P33981** | STE | 0.93 | STE | 0.85 | STE | 1.00 | 9 | STE |
| **OTHER_UHMK1/23-304 UHMK1_HUMAN UHMK1 Q8TAS1** | STE | 0.50 | STE | 0.33 | CK1 | 0.78 | 10 | TKL |
| **OTHER_ULK1/14-278 ULK1_HUMAN ULK1 O75385** | CAMK | 0.99 | CAMK | 0.50 | CAMK | 1.00 | 4 | CAMK |
| **OTHER_ULK2/7-271 ULK2_HUMAN ULK2 Q8IYT8** | CAMK | 0.99 | CAMK | 0.48 | CAMK | 1.00 | 15 | OTHER |
| **OTHER_ULK3/14-270 ULK3_HUMAN ULK3 Q6PHR2** | CAMK | 0.86 | CAMK | 0.66 | CAMK | 0.99 | 3 | CAMK |
| **OTHER_ULK4/4-280 ULK4_HUMAN ULK4 Q96C45** | STE | 0.96 | STE | 0.79 | STE | 0.98 | 10 | TKL |
| **OTHER_WEE1/299-569 WEE1_HUMAN WEE1 P30291** | CAMK | 0.97 | STE | 0.43 | TKL | 0.67 | 10 | TKL |
| **OTHER_WEE2/212-486 WEE2_HUMAN WEE2 P0C1S8** | CAMK | 0.88 | STE | 0.29 | CK1 | 0.89 | 10 | TKL |
| **OTHER_WNK1/220-479 WNK1_HUMAN WNK1 Q9H4A3** | STE | 0.87 | STE | 0.89 | STE | 0.99 | 9 | STE |
| **OTHER_WNK2/194-453 WNK2_HUMAN WNK2 Q9Y3S1** | STE | 0.90 | STE | 0.86 | STE | 0.99 | 9 | STE |
| **OTHER_WNK3/146-405 WNK3_HUMAN WNK3 Q9BYP7** | STE | 0.79 | STE | 0.87 | STE | 0.97 | 9 | STE |
| **OTHER_WNK4/173-432 WNK4_HUMAN WNK4 Q96J92** | STE | 0.93 | STE | 0.84 | STE | 1.00 | 9 | STE |
| **OTHER_RPS6KC1/337-1056 KS6C1_HUMAN RPS6KC1 Q96S38** | AGC | 1.00 | NEK | 0.37 | CMGC | 1.00 | 23 | OTHER |

**SI Table 6.** Parameters tested for each classifier. The values in bold correspond to the ones used for the final models

| **Classifiers** | **Hyperparameters** | **Values** |
| --- | --- | --- |
| Logistic Regression | *C* | 0.001, 0.01, 0.1, **1** |
| Random Forest | *n_estimators*  *max_depth* | 100, 200, **300**  None, 10, 20, **30** |
| Gaussian Naïve Bayes | *var_smoothing* | **1e-9**, 1e-8, 1e-7, 1e-6, 1e-5, 1e-4, 1e-3, 1e-2, 1e-1 |

**SI Table 7. Predictor performance.** Detailed performance of the supervised model for each fold in 10-fold classification.

|  | **LR Accuracy** | **LR MCC** | **RF Accuracy** | **RF MCC** | **GS Accuracy** | **GS MCC** |
| --- | --- | --- | --- | --- | --- | --- |
| Fold 0 | 0.97 | 0.97 | 0.95 | 0.94 | 0.95 | 0.94 |
| Fold 1 | 0.97 | 0.97 | 0.97 | 0.97 | 0.97 | 0.97 |
| Fold 2 | 1 | 1 | 1 | 1 | 0.95 | 0.94 |
| Fold 3 | 1 | 1 | 0.95 | 0.94 | 0.93 | 0.91 |
| Fold 4 | 0.97 | 0.97 | 1 | 1 | 0.97 | 0.97 |
| Fold 5 | 0.95 | 0.94 | 0.97 | 0.97 | 0.86 | 0.82 |
| Fold 6 | 0.95 | 0.94 | 0.95 | 0.94 | 0.97 | 0.97 |
| Fold 7 | 0.97 | 0.97 | 0.95 | 0.94 | 0.95 | 0.94 |
| Fold 8 | 0.95 | 0.94 | 0.93 | 0.91 | 0.93 | 0.91 |
| Fold 9 | 0.93 | 0.91 | 0.95 | 0.94 | 0.86 | 0.83 |
| Mean | 0.96 | 0.96 | 0.96 | 0.95 | 0.93 | 0.92 |

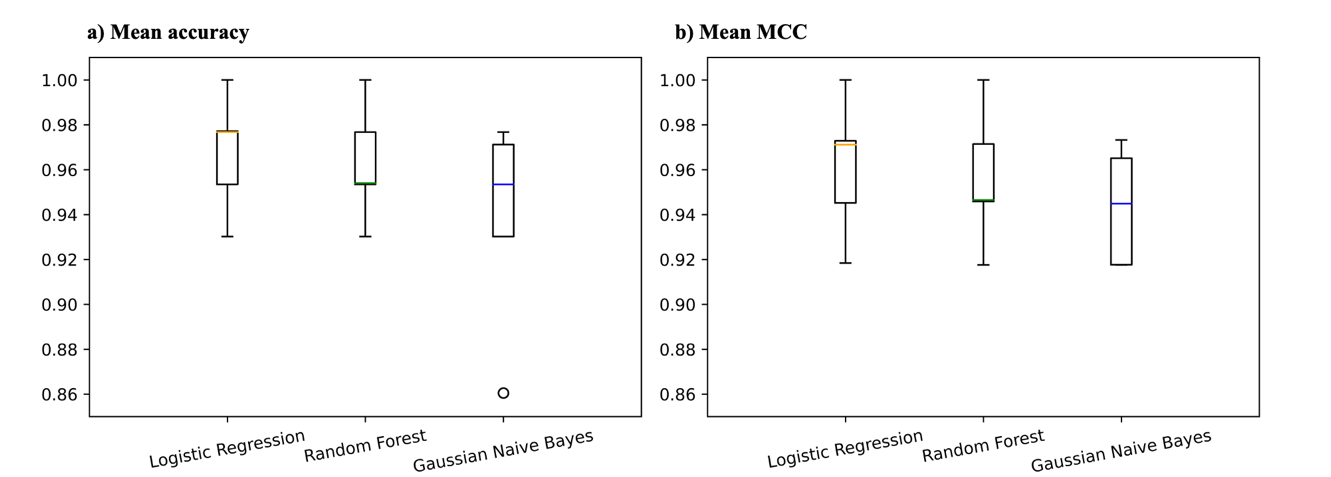

**SI Figure 8. Model performance boxplot. a)** mean Accuracy for different models across folds. **b)** mean MCC for different models across folds.

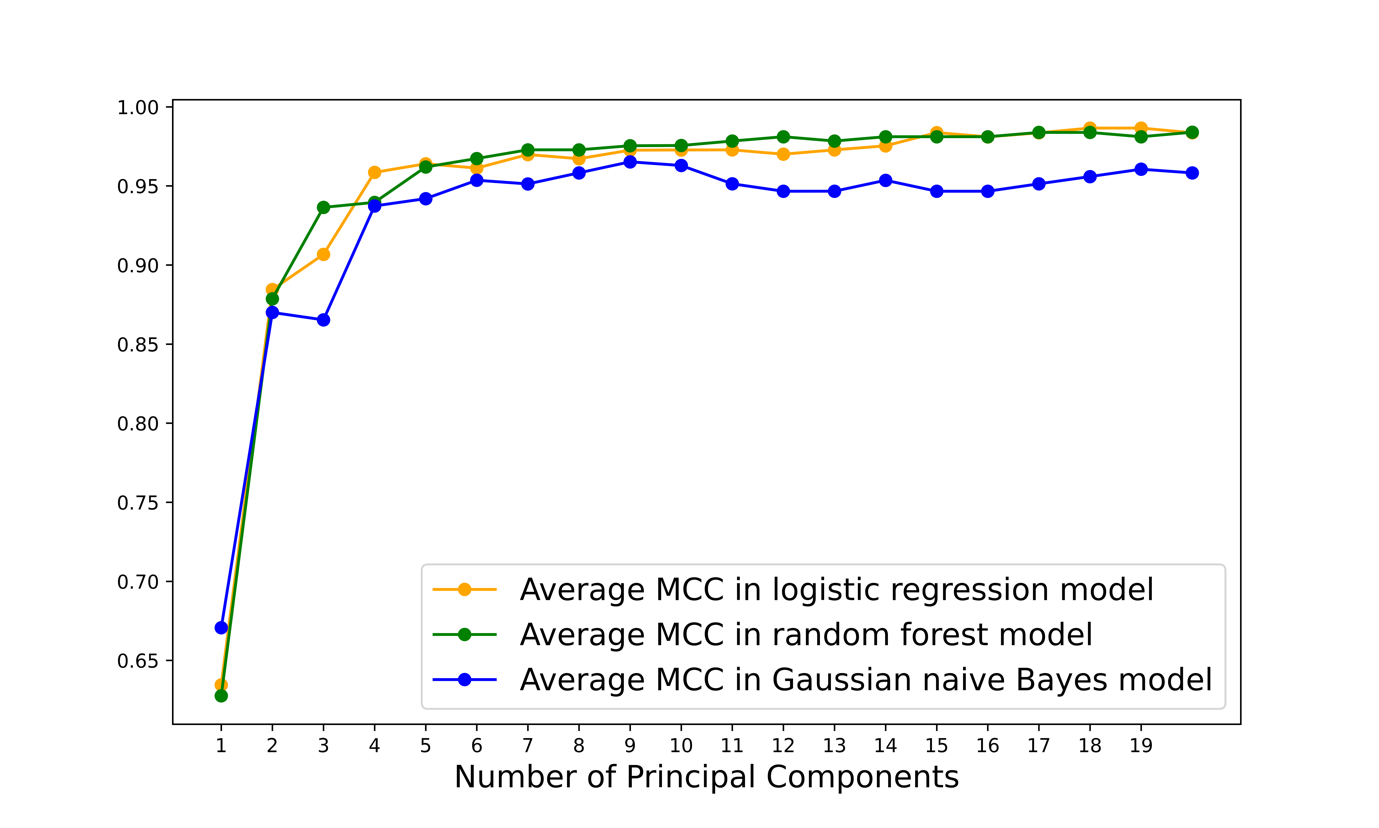

**SI Figure 9.** Average MCC of the best performing hyperparameters of each model containing different PC numbers (x-axis)

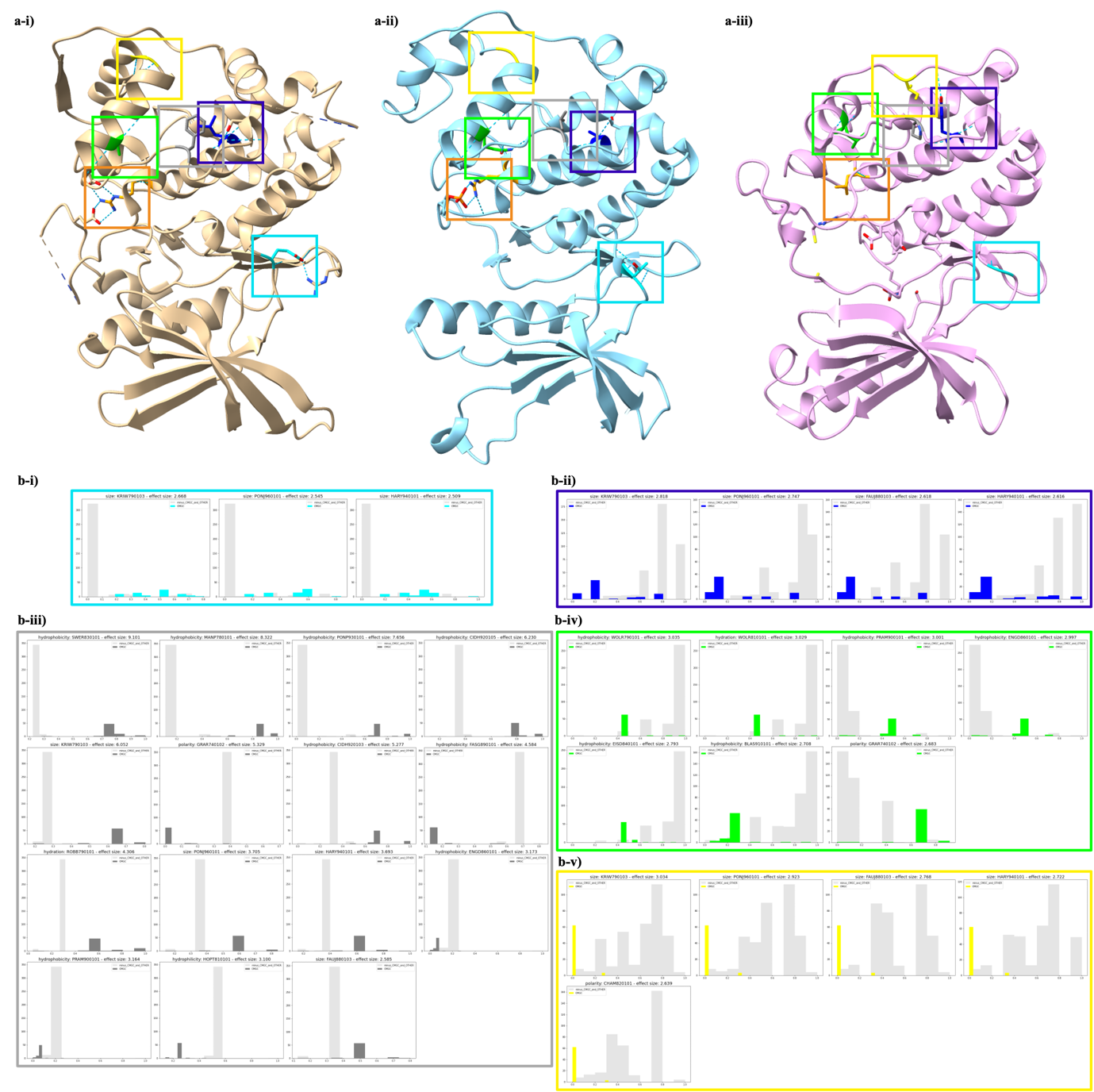

**SI Figure 10.** Highlighting more significant CMGC positions and a side by side comparison of CDC7 (unlabeled kinase classified as CMGC by our models) to CMGC and AGC. **a-i)** 4F9B[42] of unlabeled kinase CDC7 unanimously predicted in the CMGC class by all models**. a-ii)** 4MYG is phosphorylated structure of a CMGC kinase**. a-iii)** 6HHJ[41] labeled as AGC. **b-i)** Position 44, **b-ii)** Position 1969, **b-iii)** Position 1994, **b-iv)** Position 2053, and **b-v)** Position 2064

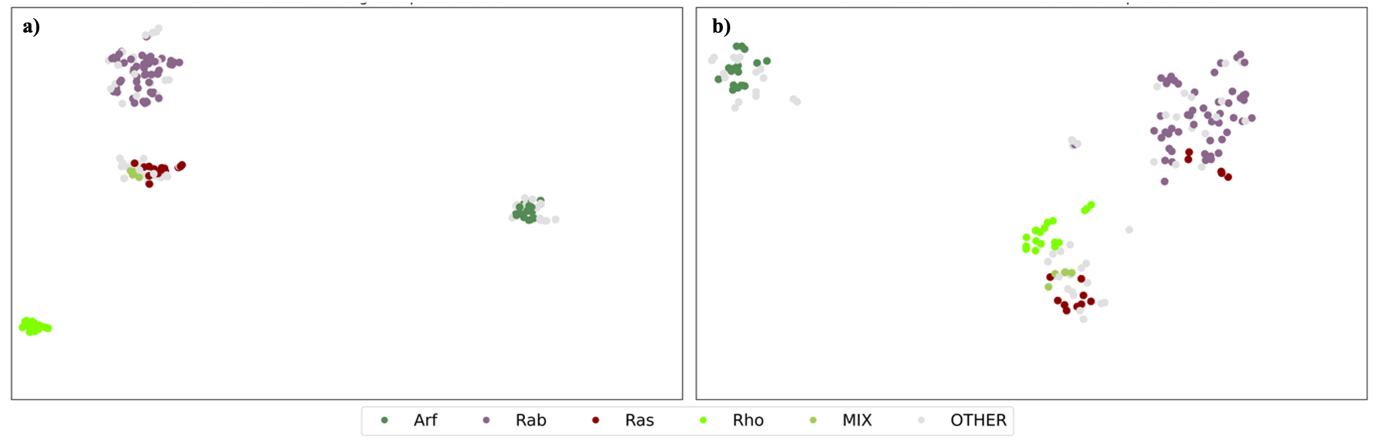

**SI Figure 11. Our method applied to small GTPase family. UMAP visualization color coded based on main class labels**. **a)** UMAP visualization of the original small GTPase representation. **b)** UMAP visualization of the PCA-transformed small GTPase representations. Importantly, representation vectors do not contain any information about the classes.

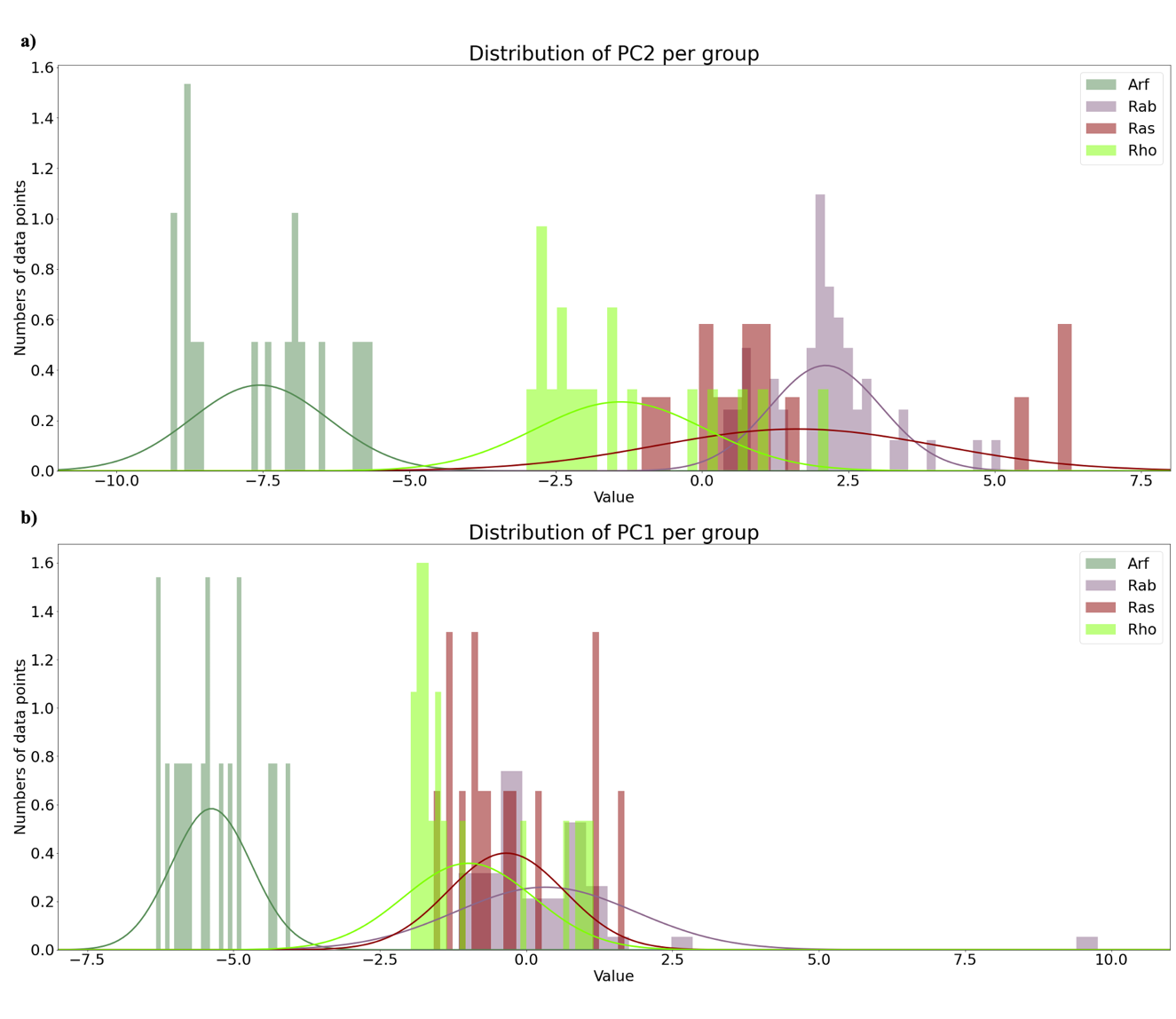

**SI Figure 12. Two first PC distributions of the small GTPase analysis with our method. a)** PC1 and **b)** PC2 distributions, color coded based on assigned classes. X-axis represents the PC values and the y-axis the number of data points, also known as the Probability Density.
